# MorphNet Predicts Cell Morphology from Single-Cell Gene Expression

**DOI:** 10.1101/2022.10.21.513201

**Authors:** Hojae Lee, Joshua D. Welch

**Affiliations:** Department of Electrical and Computer Engineering, University of Michigan, Ann Arbor, MI; Computer Science and Engineering, University of Michigan, Ann Arbor, MI; Department of Computational Medicine and Bioinformatics, University of Michigan, Ann Arbor, MI

**Keywords:** Cell Morphology, Spatial Transcriptomics, Single Cell RNA-seq, Deep Generative Model

## Abstract

Gene expression and morphology both play a key role in determining the types and functions of cells, but the relationship between molecular and morphological features is largely uncharacterized. We present MorphNet, a computational approach that can draw pictures of a cell’s morphology from its gene expression profile. Our approach leverages paired morphology and molecular data to train a neural network that can predict nuclear or whole-cell morphology from gene expression. We employ state-of-the-art data augmentation techniques that allow training using as few as 10^3^ images. We find that MorphNet can generate novel, realistic morphological images that retain the complex relationship between gene expression and cell appearance. We then train MorphNet to generate nuclear morphology from gene expression using brain-wide MERFISH data. In addition, we show that MorphNet can generate neuron morphologies with realistic axonal and dendritic structures. MorphNet generalizes to unseen brain regions, allowing prediction of neuron morphologies across the entire mouse isocortex and even non-cortical regions. We show that MorphNet performs meaningful latent space interpolation, allowing prediction of the effects of gene expression variation on morphology. Finally, we provide a web server that allows users to predict neuron morphologies for their own scRNA-seq data. MorphNet represents a powerful new approach for linking gene expression and morphology.

## Introduction

The forms and functions of cells are closely related. Historically, morphology was one of the first ways of identifying distinct types of cells, a fact reflected in descriptive names such as squamous, stellate, and dendritic cells. Cell morphology serves as a crucial marker of cellular identity across a range of biological contexts. For example, cell morphology is used in diagnosing cancer by identifying irregular or enlarged nuclear morphology^1^, monitoring metastatic events through detachment from basal membrane^2^, determining differentiation status of stem cells^3^, and elucidating neural circuits through axonal and dendritic structures. More recently, single-cell transcriptome and epigenome profiling have enabled unbiased definition of cell types using molecular features, but technological limitations make it difficult to relate molecular and morphological properties. Linking single-cell molecular profiles with morphology would provide key insights to further study important functions of cells that give rise to neural circuits, stem cell differentiation, cancer metastasis, and more.

The need for linking morphological and molecular data is particularly acute for neurons, whose complex morphologies are highly cell-type-specific, essential for their function, and difficult to assay in high-throughput fashion. The BRAIN Initiative Cell Census Network (BICCN) is currently working to generate a comprehensive brain cell atlas that characterizes the neuronal cell types in the mammalian brain^6,7^. Measuring millions of single-cell gene expression profiles has enabled discovery of new cell types, gene regulatory mechanisms, and insights into developmental dynamics and neurodegenerative diseases^8,9^. However, a neuron’s transcriptome is just one of many facets that make up its identity, and other modalities such as morphology and electrophysiology are needed to create a taxonomy of cell types and to understand properties of neural circuits.

Recent advancements in spatial transcriptomics allow measurement of gene expression between thousands to millions of single cells while retaining tissue coordinates. Because spatial transcriptomic approaches sample molecules from cells in their native tissue context, some protocols allow for paired measurement of morphology and gene expression from the same single cell. For example, the spatial transcriptomic methods MERFISH^10–13^, seqFISH^14–16^, osmFISH^17^, and Seq-Scope^18,19^ all include an imaging step that gives morphological images for individual cells. Additionally, Patch-seq allows for simultaneous profiling of transcriptomics, electrophysiology, and morphology across thousands of neurons^20–23^.

Paired morphological and molecular data from the same cells provides a unique opportunity to explore the connection among these modalities, but existing studies have not used these data types to generate new morphologies. Tripathy *et al*. used statistical methods to infer cellular features from single-cell gene expression and identified 420 genes whose expression level correlated with electrophysiological features^24^. Bomkamp *et al*. similarly identified genes whose expression levels relate to electrical and morphological properties for excitatory and inhibitory neuron cell types using Patch-seq datasets^25^. Bao *et al*. developed multi-modal structured embedding (MUSE), which uses deep learning to learn a low dimensional joint representation from both gene expression and morphology from spatial transcriptomics datasets^26^. Monjo et al. developed a deep learning model for Spatial gene Clusters and Expression (DeepSpaCE), which predicts spatial-transcriptome profiles from H&E-stained images^27^.

### However, to our knowledge no existing approaches allow the prediction of a cell’s morphology from its gene expression

We present MorphNet, a computational approach that can generate possible morphologies from the gene expression profile of a cell. Our approach leverages paired morphology and molecular data^20,21^ to train state-of-the-art deep generative models^28, 31^ that can predict single-cell nuclear or whole-cell morphology given gene expression data. To overcome the challenge of training a deep learning model using limited data such as from labor intensive experiments such as Patch-seq^21,32^, we employed state-of-the-art data augmentation techniques. We find that MorphNet can generate realistic, novel morphological images that retain the complex relationship between gene expression and cell appearance. In addition, we show that MorphNet can generalize to new contexts and augment single-cell RNA-seq data by predicting morphology. We then use MorphNet to predict morphologies for neurons from the mouse isocortex and bed nucleus. Finally, we show that MorphNet performs meaningful latent space interpolation, allowing prediction of the effects of gene expression variation on morphology.

## Results

### Overview of MorphNet model

MorphNet is a deep generative model^30,31,33,34^ that predicts the morphology of a cell from its gene expression profile. Because many factors shape morphology, knowing the gene expression of a cell constrains but likely does not uniquely determine its morphology. Thus, we formulate this problem as learning to sample from a conditional distribution of morphologies given gene expression. MorphNet learns the distribution of cell morphology conditioned on gene expression by combining two types of neural networks: a variational autoencoder (VAE)^28,29,35^ that encodes each single-cell gene expression vector into a low-dimensional representation and a generative adversarial network (GAN)^30,31,34^ that generates a distribution of possible morphology images from the gene expression representation (Figure 1**a**). The VAE and GAN are separately trained, enabling rapid and stable training. We previously showed that combining VAEs and GANs in this fashion inherits the best properties of each type of network, allowing the learning of semantically meaningful latent representations through VAEs while also generating realistic high-dimensional samples using GANs^36^.

**Figure 1.**
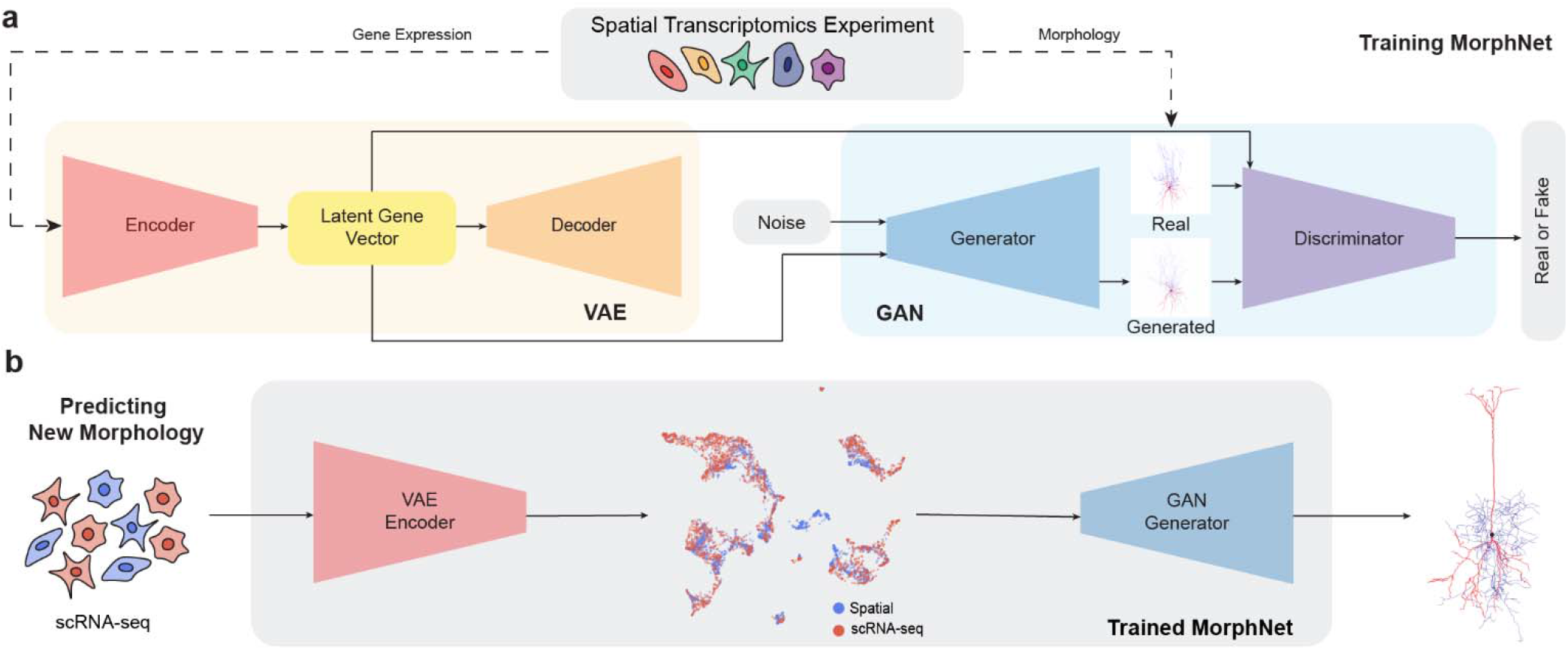
Schematic of MorphNet for predicting cell morphology from gene expression. **a.** During training, the VAE of MorphNet learns to encode single-cell gene expression into a low-dimensional gene vector. The GAN generator then takes pairs of latent gene vectors and images and learns to generate images that the discriminator cannot distinguish from the training data. **b.** MorphNet can then be used to predict new morphologies from experiments with no corresponding morphologies (such as scRNA-seq) by encoding the scRNA-seq data into the same latent space as the training data.

The encoder of the VAE model^28^ of MorphNet learns a nonlinear mapping from high-dimensional gene expression space to low-dimensional latent gene space. We use a VAE with fully-connected (multilayer perceptron) layers and a negative binomial likelihood^29,35^. The VAE is a Bayesian model that infers a distributional estimate for the latent representation of each cell’s gene expression, allowing many representations to be sampled for each single cell. The generator of the GAN^30,37^ model then takes the sampled latent gene expression vector as conditional information, which is concatenated with a small noise vector to produce a range of plausible morphological images for a given gene expression profile. We adapted the StyleGAN2-ADA architecture^31,33,34^, a state-of-the-art model that uses 2D convolutional layers to generate realistic, high-resolution images. In addition, we incorporate adaptive discriminator augmentation (ADA), which uses invertible image transformations to perform data augmentation. This allows allowing GAN training with only ~10^3^ images—many fewer than the 10^5^-10^6^ normally required for GAN training^31^. The probabilistic nature of both VAE and GAN models the many-to-many relationship between gene expression and morphology. Rather than a single gene expression profile uniquely determining a single morphology, MorphNet flexibly captures the complex relationship between a single-cell gene expression profile and the many possible corresponding morphologies.

MorphNet is trained on paired single-cell gene expression and morphological images from spatial transcriptomics experiments, such as MERFISH^11,13,38,39^ or Patch-seq^20,22,40,41^. We first train the VAE to encode single-cell gene expression into the latent space and decode the original input from the low-dimensional representation^28,29^. Then we train the GAN using pairs of gene expression representations and their corresponding real images. During training, the GAN generator learns to output images that look sufficiently similar to the real images to fool the discriminator network while preserving the correspondence relationship between each gene expression profile and its real image (Figure 1**a**). Once trained, MorphNet can output a distribution of morphological images conditioned on any latent gene expression vector, allowing it to generalize beyond the training data and predict morphologies from unseen gene expression profiles. To predict morphological images of scRNA-seq datasets that lack morphological data, we first combine the datasets with spatial transcriptomic datasets to obtain a shared latent gene space. After training the GAN on the shared latent gene expression with existing spatial transcriptomic training data, MorphNet can infer new morphological images for scRNA-seq cells in the shared latent space (Figure 1**b**).

### MorphNet generates realistic nuclear morphology images from gene expression

We first evaluated MorphNet on data from the MERFISH protocol^39^, which uses multiplexed fluorescent *in situ* hybridization to measure gene expression with subcellular spatial resolution^11,13,38^. Most MERFISH datasets also include DAPI fluorescent imaging, which highlights nuclear morphology and is used to aid cell segmentation. We reasoned that the DAPI channel gives potentially useful morphology data that can be paired with the gene expression measurements from the same cell. We focused in particular on the Vizgen Mouse Brain Receptor Map dataset, which contains measurements for 483 genes from 734,696 cells across 9 coronal slices from the adult mouse brain^39^. We segmented the 100-nm resolution DAPI images using the provided single-cell boundaries. Note that we used the cell boundaries, which subsume the nucleus; thus, the boundary of the nucleus is determined by the measured DAPI signal rather than a computational segmentation algorithm. To ensure uniform image sizes, we centered and padded each nuclear image to 256 ×256 pixels.

MorphNet generated images reflect the conditional information from gene expression. To do this, we first trained a deep neural network classifier^43^ to recognize the transcriptional cell type of a cell from its nuclear morphology using the real image data. Then, we evaluated the generated images using the trained classifier for which the true transcriptional labels are known. If MorphNet respects the relationship between gene expression and morphology, the classifier performance will be similar when evaluated on either real or generated images.

We also implemented a baseline model to compare with MorphNet (**Supplementary Figure 1**). The baseline model uses a convolutional neural network architecture^43–47^, and consists of transpose convolutional layers and convolutional blocks inspired by U-Net decoders^48^. We trained the baseline model to predict images from a latent gene expression representation using mean squared error (MSE) loss. We then compared the images generated by the baseline model with images generated by MorphNet, which was trained using adversarial loss (Figure 2**a**).

**Figure 2.**
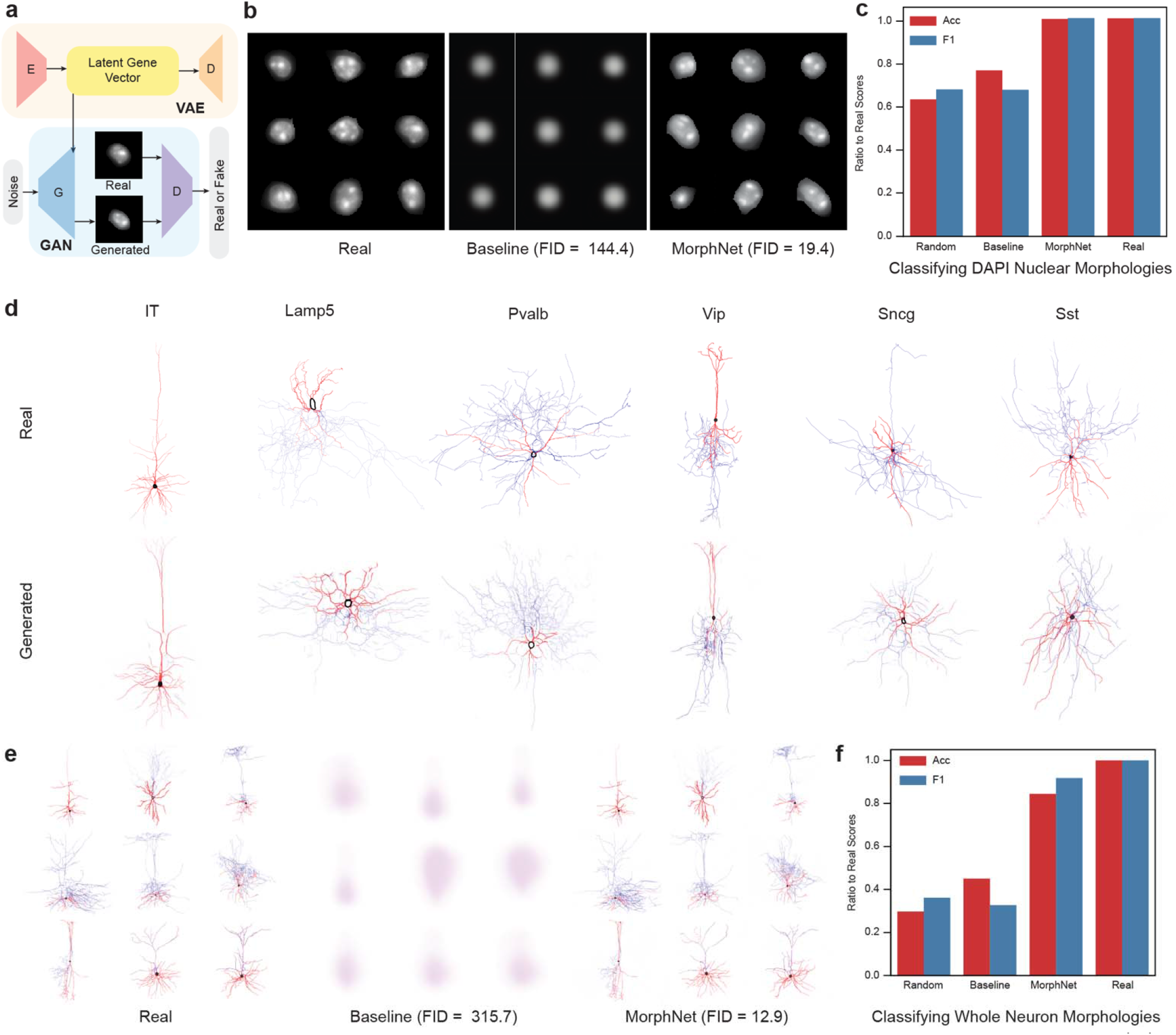
MorphNet generates realistic morphology images from single-cell gene expression. a. Training scheme of MorphNet on Vizgen MERFISH dataset. **b.** Comparison between real single-cell nucleus images and generated morphological images from baseline and MorphNet. MorphNet can generate complex, realistic nuclear shapes, while the baseline model cannot. c. Relative accuracy and F1 scores from classifying real or qenerated DAPI Nuclei images as from neuron or non-neuronal cell type. d. Comparison between real neuron morphologies and those generated from a holdout set using MorphNet across six cell types present in the Patchseq dataset. e. Comparison between real Patch-seq images and generated whole neuron morphologies from baseline and MorphNet. f. Relative accuracy and F1 scores from classifying the transcriptional cell type of real or generated neuron morphologies using a classifier trained on the real images. Note that the accuracy and F1 score are reported relative to the accuracy of the classifier evaluated on the real data.

MorphNet generates highly realistic nuclear morphology images from the MERFISH data. The generated images are qualitatively very similar to the real images and reflect nuances of complex shape and fluorescence intensity patterns (Figure 2**b**). In contrast, the generated images from the baseline model match the real images with respect to overall fluorescence intensity and total nuclear area but fail to capture more complex nuclear shapes, often producing approximately circular single-nucleus images. The FID metrics also indicate that MorphNet generates much more realistic images than the baseline: MorphNet achieves an FID of 19.4, while the FID for the baseline model is 144.4 (Figure 2**b**). The relatively poor performance of the baseline model is likely because the assumption that each gene expression profile uniquely determines a single morphology image is invalid. Instead, gene expression and morphology have a complex, many-to-many relationship in which gene expression profiles constrain but do not uniquely determine the morphological images.

MorphNet also generates images that retain the relationship between gene expression and morphology. Although nuclear morphology is not as cell-type-specific as whole-cell morphology, our trained classifier was able to distinguish neuronal nuclei from non-neuronal nuclei with 80.4% accuracy and 83.5% F1 score. Thus, the morphology of the real images does reflect the transcriptional cell type to some detectable degree. When evaluated on MorphNet images, the classifier achieved an accuracy of 99.7% and F1 score of 100% relative to the real data (Figure 2**c**), indicating that MorphNet respects the constraint of the gene expression information when generating morphologies. In comparison, classification of generated images from the baseline models performs similarly to that of a random classifier (**Supplementary Figure 2**), indicating the lack of encoded cell type information in the generated images.

We trained MorphNet on an additional spatial transcriptomic data type called cyclic-ouroboros smFISH, or osmFISH (**Supplementary Figure 3**). We found that MorphNet again produced highly realistic images of nuclear morphology from DAPI staining (FID = 6.89 for training set; FID = 8.37 for testing set). However, the osmFISH morphology data was lower resolution than that of MERFISH data, and a classifier trained to recognize transcriptional cell types from the real morphologies did not outperform a random baseline. This indicated that the real morphology images lacked information about the transcriptomic cell type, so we did not analyze the osmFISH dataset further.

### MorphNet generates realistic whole-neuron morphology images from gene expression

We next applied MorphNet to generate whole neuron morphological images using Patch-seq data^21,32^. The Patch-seq protocol measures transcriptomic, electrophysiological, and morphological properties from the same cells^41^. Whole-neuron morphology images from Patch-seq are significantly more complex than nuclear morphology images, containing intricate dendrite and axon branching structures. In addition, the number of cells is limited due to the laborious manual processes required for patch clamping and morphological reconstruction. However, neuron morphology is much more cell-type-specific than nuclear morphology, and the ability to predict neuron morphology from gene expression would be a powerful tool for understanding the functional implications of molecular variation.

To train MorphNet to generate whole neuron morphologies, we combined two Patch-seq datasets from the mouse visual cortex and mouse motor cortex^21,32^. The combined datasets include 5,764 cells with gene expression measurements. We annotated each of the cells according to six broad transcriptional cell types: intratelencephalic-projecting excitatory neurons (IT), interneurons expressing vasointestinal peptide (*Vip*), interneurons expressing *Lamp5*, interneurons expressing parvalbumin (*Pvalb*), interneurons expressing somatostatin (*Sst*), and interneurons expressing *Sncg*. As with the MERFISH data, we first trained a VAE model with negative binomial likelihood to encode Patch-seq gene expression measurements into 10-dimensional latent gene representations. To obtain whole neuron morphological images, we projected the reconstructed neuron morphologies onto the *xy*-plane and colored the pixels by neuronal component (dendrite = red; axon = blue; soma = black; **Supplementary Figure 4**). We then split the dataset into training set (90%) and testing set (10%) stratified by cell type and trained MorphNet on the paired image and gene expression data.

Despite the relatively small number of morphology images available for training, MorphNet was able to generate highly realistic neuron morphologies that reflect the provided gene expression data (Figure 2**c**). We experimented with different GAN architectures^31,49^ (**Supplementary Figure 5**), and found that StyleGAN2^31^ is able to capture the complex morphologies of whole neurons. We further tested a data augmentation technique called adaptive discriminator augmentation (ADA)^31^ and found it to significantly reduce overfitting in the discriminator network, improving MorphNet’s generalizability to unseen data (**Supplementary Figure 6**). We compared the true and generated morphology images for held-out cells; the network had not seen either the gene expression profile or the corresponding morphology. Note that we also did not provide any cell type labels to MorphNet during training. Nevertheless, when given new Patch-seq gene expression profiles, MorphNet was able to generate realistic morphologies that were strikingly similar to the true corresponding morphologies and retained the distinctive characteristics of the six transcriptional cell types (FID = 12.90 for training set; FID = 15.82 for testing set). This indicates that MorphNet successfully generalized to the test set despite the limited training data.

The baseline model achieved an FID score of 315.7 compared to the FID score of 15.82 from MorphNet (recall lower FID is better). Visually, the images generated by MorphNet are nearly indistinguishable from real neuron morphologies (Figure 2**d**). As with the MERFISH data, the baseline model generates highly blurred images that reflected some characteristics of the neuron color and shape, but do not resemble neurons at all. We also tested whether the generated images contain cell-type specific information by training a ResNet-50 classifier^43^ to recognize the transcriptional cell type (IT, *Lamp5, Vip, Pvalb, Sst*, and *Sncg*) from the generated morphology. Because neuron morphology is highly cell-type-specific, we were able to classify the generated morphology images according to transcriptional cell type with 73% accuracy and 60% F1 score. The same classifier evaluated on the images generated by MorphNet achieved 84.4% accuracy and 91.8% F1 score relative to the classifier performance on the real images. A baseline classifier that randomly guesses the cell type with probability equal to the class proportions performed significantly worse, with a relative accuracy score of only 29.7% and relative F1 score of 36.2%. In addition, the images generated by the baseline model scored a relative accuracy score of 45.0% and a relative F1 score of 32.7%, indicating that the baseline model does not generate recognizable examples of the morphologies from each transcriptional cell type (**Supplementary Figure 7**). This indicates that, overall, the generated images retain morphological differences among transcriptional cell types and accurately reflect the complex relationship between gene expression and morphology (Figure 2**e**).

Recent papers have shown that trained GAN generators can be used to discover nonlinear directions of variation in data space62. In the case of MorphNet, such directions would correspond to the dominant ways in which morphologies vary among cells.are among the primary sources of variation in MorphNet generation results, while for neuron morphologies, the degree of axon and dendrite branching are the dominant morphological axes.

### MorphNet predicts morphology from unseen single-cell RNA-seq profiles

The ability of MorphNet to predict morphology from gene expression raises the exciting possibility of augmenting sc RNA-seq datasets with inferred morphologies. To investigate this further, we tested whether MorphNet can generate realistic whole neuron morphology for scRNA-seq datasets that have no experimentally measured morphology information. We retrained the MorphNet VAE to embed both Patch-seq and scRNA-seq data in the same latent space. Then we retrained the GAN using the joint latent space for both data types and the paired images for the Patch-seq dataset. Finally, we predicted morphologies for the scRNA-seq profiles by passing them through the VAE encoder and using the latent representation as a condition for the GAN generator (Figure 3**a**). We used this same training scheme for three different scRNA-seq datasets from different regions of the adult mouse brain.

**Figure 3.**
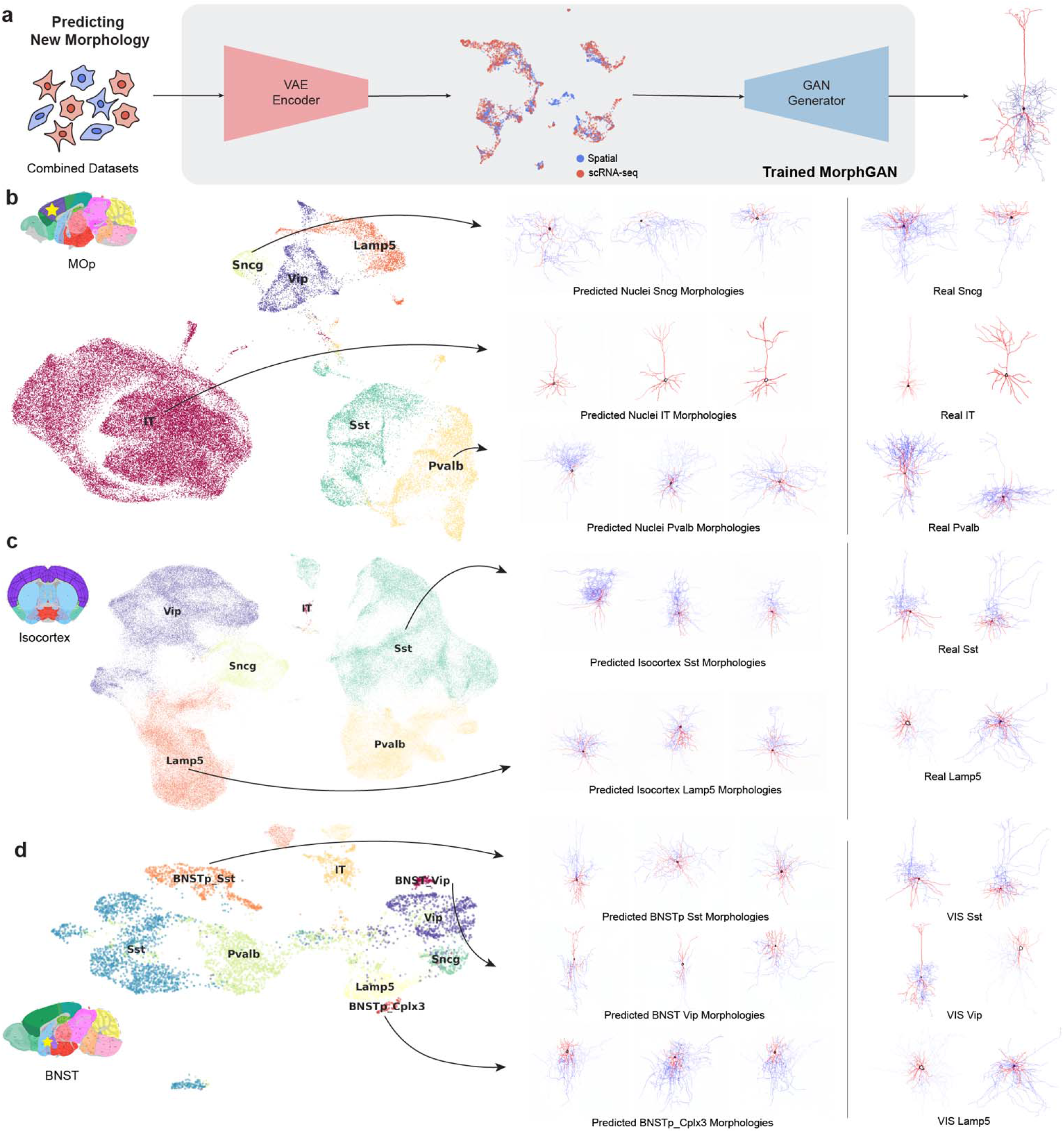
MorphNet predicts morphologies for scRNA-seq datasets. **a.** Schematic of predicting new neuron morphologies for scRNA-seq datasets with no ground truth morphologies. **b.** UMAP plot of latent gene expression space from Patch-seq + MOp colored by cell type. Predicted whole neuron morphologies for *Sncg*, IT, and *Pvalb* cell types from MOP dataset on left and samples of ground truth images from Patch-seq on right. **c.** UMAP plot of latent gene expression space from Patch-seq + Isocortex colored by cell type. Predicted whole neuron morphologies for *Sst* and *Lamp5* cell types on left and samples of ground truth images from Patch-seq on right. **d.** UMAP plot of latent gene expression space from Patch-seq + BNST colored by cell type. Predicted whole neuron morphologies for cells from BNST dataset on left and real morphologies from visual cortex (VIS) Patch-seq data on right.

We first tested MorphNet on a dataset containing 215,823 single-nucleus RNA-seq (snRNA-seq) profiles from the primary motor cortex (MOp) of the adult mouse brain^50^. The motor cortex is near the visual cortex and contains transcriptionally similar cell types, and the Patch-seq training data already contains cells from the motor cortex^51^. Thus, we expect that MorphNet should be able to generate reasonable morphological predictions for snRNA-seq data from MOp. Prior to obtaining gene expression encodings, we filtered the MOp dataset to only cell types that exist in the Patch-seq dataset. Figure 3**b** shows the UMAP plot of the latent gene expression space for the combined Patch-seq and MOp datasets. As shown in **Table 1**, using combined latent gene expression space from Patch-seq and MOp achieved an FID score of 14.68 on the Patch-seq dataset alone and FID score of 14.83 on the MOp dataset alone. Figure 3**b** shows predicted neuron morphologies from *Sncg*, IT, and *Pvalb* cell types from the MOp dataset as well as sample images from corresponding cell types from Patch-seq. **Supplementary Figure 8** shows uncurated generated images of all cell types in the MOp dataset. The images generated from snRNA-seq profiles are highly realistic and qualitatively retain the cell-type-specific morphologies seen in the Patch-seq cells, with a relative accuracy score of 57.52% and relative F1 score of 64.25%. Thus, MorphNet successfully predicts morphologies even for snRNA-seq datasets with no morphological information.

**Table 1.** Metrics on generated neuron morphologies from MorphNet trained on combined datasets.

| Datasets | FID score |  | Classification |  |
| --- | --- | --- | --- | --- |
|  | Patch-seq | scRNA-seq | Rel. Acc. | Rel. F1 |
| Patch-seq + MOp | 14.68 | 14.83 | 57.52 | 64.25 |
| Patch-seq + Isocortex | 14.64 | 32.30 | 64.01 | 71.27 |
| Patch-seq + BNST | 14.54 | 38.45 | 90.92 | 100 |

Next, we tested MorphNet on a large scRNAseq dataset from the entire Isocortex region of the adult mouse brain. We subset the Isocortex dataset to only GABAergic cells and combined with Patch-seq for a total of 176,272 cells and 7,595 genes. Even though Patch-seq data is from only visual and motor cortex, sub-regions of the Isocortex, we hypothesized that we could predict accurate morphologies for other cortical regions because cortical GABAergic neuron gene expression does not show significant regional variation^51^. Figure 3**c** shows the UMAP plot of the learned gene expression space for the combined datasets, as well as predicted neuron morphologies for cells in the Isocortex dataset. **Supplementary Figure 9** shows uncurated generated images for GABAergic cell types in the Isocortex dataset. As with the MOp, the morphologies generated from scRNA-seq profiles look realistic and retain the characteristic morphologies of different transcriptional types with FID score of 32.30 and relative accuracy of 64.01% and relative F1 score of 71.27%. This demonstrates MorphNet’s ability to generate and predict neuron morphologies for scRNAseq data across a large region of the mouse brain.

Having established that MorphNet can predict morphologies for scRNA-seq profiles from new areas of the cortex, we applied the method to a brain region outside the cortex. We previously reported diverse transcriptional subtypes of GABAergic neurons in the bed nucleus of stria terminalis (BNST), including several clusters that express some of the same key marker genes as cortical interneurons^52^. To determine which BNST cells are most similar to cortical interneurons, we integrated the MOp snRNA-seq dataset with our previous BNST snRNA-seq dataset using the LIGER^52^ algorithm. This analysis showed that the cell types in MOp and BNST are quite different overall, with very few overlapping populations (Supplementary Figure 10**a**). However, we did identify three minor populations within the BNST, constituting 1,053 cells (2.7% of all BNST cells), which aligned with cortical interneuron types (Supplementary Figure 10**b**). We used LIGER to identify *k* – 80 joint clusters and calculated the percentage of each cell type from both MOp and BNST within each cluster. While most clusters contain cells from only one region, we found that the BNST_Cplx3 cell type aligns with the Lamp5 cluster, the BNST_*Sst* cell types aligns with the *Sst* cortical type, and the BNST_*Vip* matches with the *Vip* cluster from MOp. We thus extracted only the cells from these three BNST clusters and trained a VAE model to jointly encode these cells with the Patch-seq data (**Figure 3d**). A UMAP visualization of the VAE latent space also supports the similarity of these cell types, with BNST_*Sst* positioned near *Sst*, BNST_*Vip* positioned near *Vip*, and BNST_*Cplx3* positioned near *Lamp5*. We next used MorphNet to predict morphologies for the BNST cells (**Figure 3d**). **Supplementary Figure 11** shows uncurated generated images of the BNST dataset. Remarkably, despite the significant transcriptional differences between the Patch-seq cells used in training and the BNST cells, MorphNet predicted realistic morphologies BNST snRNA-seq profiles. To our knowledge, the morphologies of these cells have not been experimentally characterized. However, some studies have predicted that BNST_Cplx3 morphology could be similar to that of the cortical neurogliaform neurons^53^. The MorphNet predictions further support this hypothesis.

### Latent space interpolation predicts morphological effects of varying gene expression

A key benefit of deep generative models is their ability to perform latent space interpolation—to predict realistic high-dimensional examples from unseen combinations of latent variables. In the context of cell morphologies, interpolation in the latent gene expression space would allow MorphNet to predict the morphological effects of new combinations of expressed genes. Such an ability could not only yield interesting predictions, but could reveal new insights into the specific ways in which gene expression influences morphology.

We first investigated the effects of linear interpolation between the latent embeddings of Patch-seq cells. To do this, we chose a starting cell (such as a *Lamp5* neuron) and an ending cell (such as a *Vip* neuron). Then we calculated new latent space positions by taking a weighted average of the start and end cell embeddings, with varying weights—a process called linear interpolation. Note that these interpolated latent space positions do not result from encoding the gene expression profile for a real cell, but instead correspond to hypothetical unobserved gene expression profiles intermediate between the start and end neurons. We then generated morphologies for the interpolated latent space positions (Figure 4**a**). Remarkably, the predicted morphologies show meaningful interpolation, with the intermediate latent space positions still producing realistic neuron morphologies that gradually transition from the morphology of the starting cell to the morphology of the end cell. We repeated this procedure 100 times and confirmed that the images generated from interpolated gene expression profiles consistently interpolated among the start and end cell morphologies, including when varying the transcriptional types of the start and end cells. A classifier trained on the real images confirms this qualitative result, with gradually decreasing classification probabilities for the starting cell type *Lamp5* (70.46% ± 6.98%, 45.71% ± 7.44%, 5.92% ± 2.65%, 2.74% ± 1.67%, and 1.49% ± 1.75%) and gradually increasing probabilities of the ending cell type *Vip* (5.50% ± 3.75%, 7.11% ± 3.74%, 37.43% ± 7.68%, 60.22% ± 7.50%, and 84.75% ± 4.73%). These numbers quantitatively confirm the morphological changes reflected in the predicted images. We also show a sampling of uncurated examples in Supplementary Figure 12 (linear interpolation between *Lamp5* and *Vip*) and Supplementary Figure 13 (linear interpolation between *Pvalb* to *Sst*).

**Figure 4.**
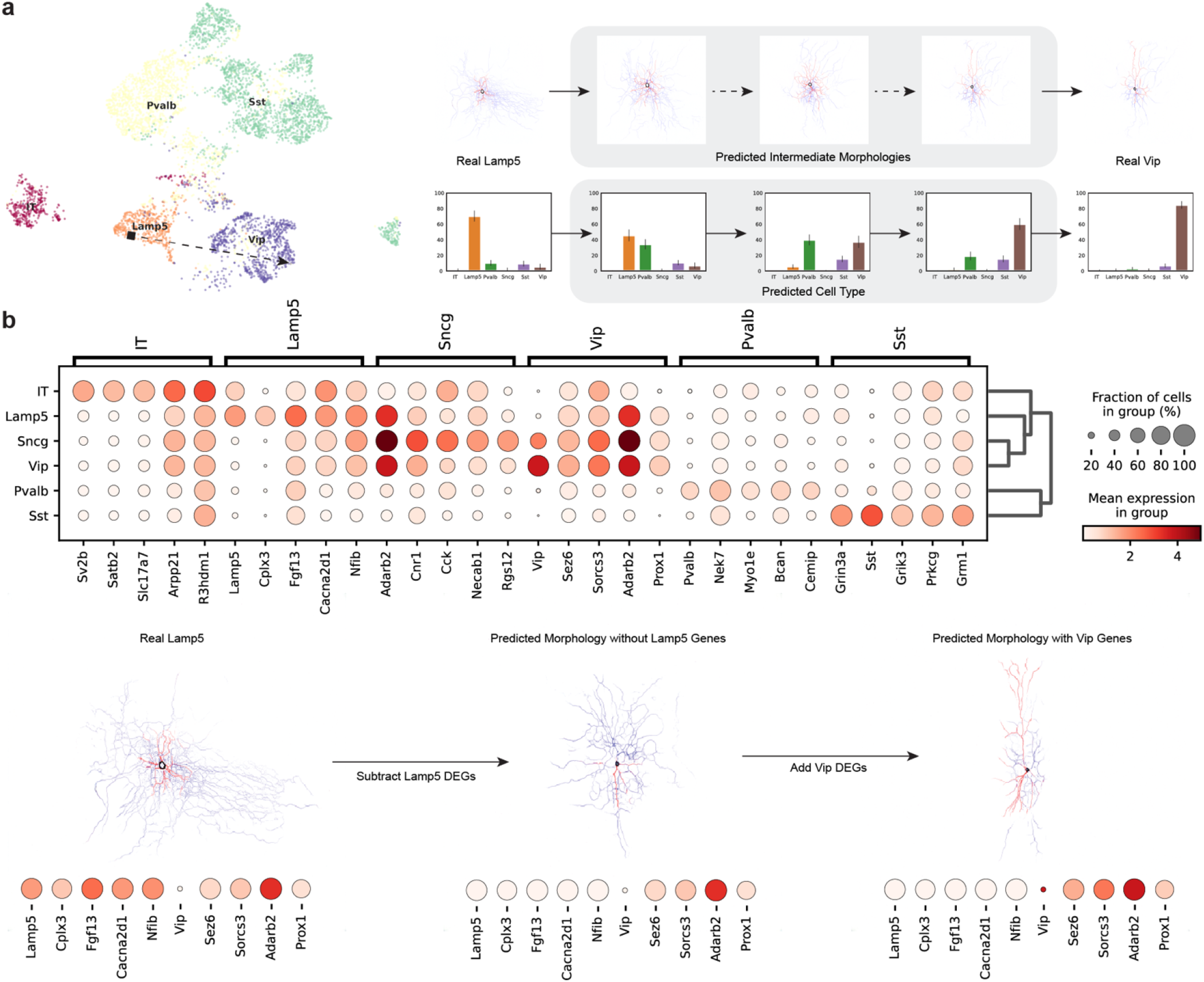
**a.** Results from linear interpolation in 10-dimensional latent gene space. Predicted morphologies between real Lamp5 and real Vip cell by MorphNet, and corresponding cell type probabilities. **b.** Dot plot of top five differentially expressed genes (DEGs) for each cell type in Patch-seq dataset. Predicted morphologies for a *Lamp5* cell type without *Lamp5* DEGs and added *Vip* DEGs.

Next, we investigated how changing the expression of specific genes affects the predicted morphology. This time, rather than manipulating the latent space, we changed the high-dimensional expression profile of a cell before encoding it into the latent space. To choose genes most likely to have a large effect, we used the Wilcoxon rank-sum test to identify differentially expressed genes (DEGs) for each transcriptional cell type (Figure 4**b**). Then, we created new gene expression profiles by zeroing out the expression of DEGs in the starting cell type (e.g., *Lamp 5*). We then set the expression of DEGs for a different target cell type (e.g., *Vip*) to an arbitrary high value, encoded these synthetic gene expression profiles into the latent space using the VAE and generated morphologies for them. The resulting morphologies are still highly realistic and qualitatively reflect the changes in gene expression: subtracting the DEG expression from the starting cell type removes much of the characteristic morphology of this cell type, while adding the DEG expression from the target cell type results in a morphology resembling this type.

**Table 2.** Number of Differentially Expressed Genes Identified per Cell Type.

|  | IT | Lamp5 | Pvalb | Sncg | Sst | Vip |
| --- | --- | --- | --- | --- | --- | --- |
| # of DEGs | 753 | 261 | 460 | 306 | 293 | 275 |

## Discussion

MorphNet is a probabilistic deep generative model that can predict new morphological images (single-cell nuclear or whole-cell) given gene expression. We showed that MorphNet is a powerful yet flexible model that accurately captures the many-to-many relationship between gene expression and morphology using MERFISH and Patch-seq datasets. In addition, we tested MorphNet on three external unimodal single-cell RNA-seq datasets (MOp, Isocortex, and BNST) which have no corresponding morphological information. This demonstrates the usefulness of MorphNet to augment existing single-cell RNA-seq databases with rich morphological information, which is crucial especially in characterizing neurons and creating a comprehensive parts list of the brain. We release a simple web tool for users to upload scRNA-seq data and obtain predicted morphologies from MorphNet (**Supplementary Figure 14**).

A limitation of MorphNet is that quality of images is somewhat dependent on whether the external dataset overlaps well with existing spatial transcriptomics data. While this is less of an issue for techniques like MERFISH, which has been scaled to measure nearly millions of cells across the entire brain, it is much more challenging for Patch-seq, which has been difficult to scale due to its labor-intensive nature. As new methods develop to automate profiling of single-cells, MorphNet will learn to generate higher quality morphological images. In addition, future studies may improve quality of out-of-distribution generation, which is an active area of study in deep learning. Given its scope, flexibility, and extensibility, we anticipate that MorphNet will be a valuable tool for characterizing the link between molecular and morphological features of cells.

## Methods

### Spatial Transcriptomics Datasets

Training MorphNet requires paired single-cell gene expression morphological data. For predicting single-cell nucleus images, we used data from the Vizgen MERFISH Mouse Receptor Map (a dataset we refer to as MERFISH for short)^39^ and osmFISH^17^. For predicting single-cell neuron morphology images, we used data generated using Patch-seq^20,22,40,41^.

#### MERFISH

Vizgen’s MERFISH Mouse Brain Receptor Map is one of the largest, publicly-available single-cell spatial transcriptomics datasets consisting of 483 gene expression measurements from 734,696 cells across nine coronal slices (three full coronal slices and three biological replicates per slice) from the adult mouse brain^39^. In addition, the raw MERFISH dataset provides single-cell segmentation coordinates and 100-nm resolution DAPI images for each coronal slice^39^. To prepare the single-cell nucleus images, we applied the single-cell segmentation coordinates to from the center z-plane image (i.e. fourth z-plane out of seven z-stacked images) from each coronal slice image. Then, we centered and padded each single-cell nucleus image to a final grayscale image of size 256 pixels-by-256 pixels. For the MERFISH transcriptomics dataset, we followed a simple preprocessing procedure using the scanpy package^54^. Briefly, we first filtered out genes that are expressed in less than three cells, normalized the library size to 10^4^, and took the log transform of each gene expression value.

#### osmFISH

osmFISH is a spatial transcriptomics method which stands for nonbarcoded and unamplified cyclic-ouroboros smFISH method^17^. The osmFISH transcriptomics dataset consists of measurements of 33 genes from 6,471 cells, as well as a stitched DAPI image from the mouse somatosensory cortex^17^. The dataset provides single-cell segmentation masks derived using the watershed algorithm applied to PolyT-stained images (Supplementary Figure 3 **a**). After cropping from DAPI images using the PolyT-derived segmentation masks, we centered and padded each single-cell nucleus image to a final grayscale image of size 400 pixels-by-400 pixels. We similarly processed the osmFISH transcriptomics data as we did on the MERFISH dataset but did not see a clear clusters by cell type (Supplementary Figure 3), likely due to limited number of genes measured across the mouse somatosensory cortex.

#### Patch-seq

Patch-seq is an experimental method that can simultaneously profile single neuron gene expression, electrophysiology, and morphology^40,41^. As the name suggests, Patch-seq uses patch clamp recordings to measure electrophysiological properties of single neurons^40^. During recording, neurons can be filled with biocytin to reconstruct detailed dendritic and axonal morphologies^41^. After recordings, the cytosol and nucleus are aspirated and collected for RNA sequencing^20,22,40,41^. In our work, we analyzed two of the largest publicly available Patch-seq datasets, one from the mouse motor cortex^20^ and other from the mouse visual cortex^22^.

We identified a total of 5,764 cells (1,329 cells from motor cortex, 4,435 cells from visual cortex) with gene expression data, of which 1,220 cells (646 cells from motor cortex, 574 cells from visual cortex) had whole neuron morphology reconstructions. Morphological data for both datasets were given in SWC files, which contain 3D coordinates of neuronal compartments and their type (soma, axon, basal dendrite, or apical dendrite)^55,56^. We used the neurom package^57^ to load, process, and project the 3D morphological data into the *xy*-plane, as well as color each pixel according to its neuronal compartment type (see Supplementary Figure 4). The *xy*-plane projection was chosen over *xz-* and *yz*-planes as they visually contained more identifiable morphological features such as long axons in IT cells and bipolarity in *Vip* cells.

To combine the two gene expression datasets, we first preprocessed each dataset to normalize the library size to 10^4^ and selected 10,000 highly variable genes using the package scanpy^54^. The datasets were then concatenated using the intersection of highly variable genes, which yielded a final Patch-seq gene expression matrix of 5,764 cells and 3,552 genes. Note that only a subset of the 5,764 cells whose gene expression were profiled also had morphological reconstructions (1,220 cells). Finally, each of the 5,764 cells were annotated with one of six cell types (IT, *Lamp5, Pvalb, Vip, Sncg*, and *Sst*), which we used as ground truth labels when classifying morphology images for cell type.

### scRNA-seq Datasets

Below we describe all sc/snRNA-seq brain datasets without morphology data. To predict neuron morphological images for these datasets, we combined the transcriptomics with that of Patch-seq using common sets of genes. We then trained the VAE to obtain the latent gene expression for each cell using the dataset name as batch key to correct for batch effects. We trained MorphNet on the new latent space with existing morphological images from Patch-seq. Finally, we fed the latent gene expression from unimodal sc/snRNA-seq datasets into the generator of MorphNet to predict new morphological images.

#### MOp

The raw MOp dataset consists of measurements of 31,053 genes from 215,823 cells from the mouse cerebellum obtained using single-nucleus RNA-seq^50^. We took 104,802 cells with a class label of either glutamatergic or GABAergic and preprocessed the dataset using scanpy package^54^. Same as the MERFISH dataset, we first filtered out genes that are expressed in less than three cells, normalized the library size to 10^4^, and took the log transform of each gene expression value. Finally, we subset the dataset to select for 10,000 highly variable genes. To obtain a shared latent space with Patch-seq dataset, we concatenated the MOp and Patch-seq dataset based on overlapping genes between the two datasets and filtered for neurons with the same cell types as in Patch-seq (i.e. Layer 2/3 IT, *Lamp5, Pvalb, Sncg, Sst*, and *Vip*). The final resulting dataset consists of 59,995 cells (54,231 cells from MOp and 5,764 cells from Patch-seq) and 2,133 genes, which we used to train the VAE of MorphNet.

#### Isocortex

The Isocortex dataset is the combination of scRNA-seq measurements across the following brain regions: visual cortex (VIS, VISp, VISl, VISm), primary motor cortex (MOp), primary somatosensory cortex (SSp), auditory cortex (AUD), prelimbic – infralimbic – orbital area (PL-ILA-ORB), retrosplenial area (RSP), entorhinal cortex (ENT), temporal association area – perihinal area – ectohinal area (Tea-PERI-ECT), anterior cingulate area (ACA), agranular insular area (AI), PAR-POST-PRE-SUB-ProS, SSs-GU-VISC-AIp, secondary somatomotor area (MOs_FRP), and posterior parietal association area (PTLp)^58^. The combined raw dataset contains 1,169,320 cells by 31,053 genes. For each of the brain regions, we removed duplicate genes and preprocessed the dataset as in MERFISH (filtering, normalizing, and log transform). We also excluded the cells from hippocampus (HIP) and filtered for 170,508 GABAergic cells. After selecting for 10,000 highly variable genes and concatenating with the Patch-seq dataset, we obtained a final cell-by-gene matrix of 176,272 cells (170,508 cells from Isocortex and 5,764 cells from Patch-seq) by 7,595 genes.

#### BNST

The bed nucleus of stria terminalis (BNST) is in the subcortical region of the brain and associated with social, stress-related, and reward behaviors^52^. The raw dataset contains 38,806 cells and 24,301 genes belonging to one of 41 clusters. Exploratory data analysis of the BNST dataset with MOp showed poor alignment using the VAE^29^ (Supplementary Figure 10**a**). To select clusters of BNST cells most similar to cortical interneurons, we used the LIGER^52^ algorithm to identify 80 joint clusters between the BNST dataset and MOp dataset. We examined each of the 80 clusters for percentage of each cell type present in the datasets. From this analysis, we identified three BNST clusters that align with the cortical cell types, i.e. BNST_*Cplx3* with the *Lamp5* cluster, BNST_*Sst* with the MOp *Sst* cluster, and BNST_*Vip* with the MOp *Vip* cluster. After subsetting for these three cell types, we preprocessed the dataset as we did for the MOp dataset (filtering, normalizing, log transform). We then selected for 10,000 highly variable genes and concatenated with the Patch-seq dataset. The final dataset contains 6,817 cells (1,053 cells from from BNST and 5,764 cells from Patch-seq) and 3,051 genes, which was used to train the VAE of MorphNet.

## Models

### Baseline

The CNN-based benchmark model is analogous to the decoder network from U-Net^48^ and schematically depicted in Supplementary Figure 1. Our benchmark model is a fully convolutional neural network consisting of transpose convolution layers^59^ to increase spatial dimension and two blocks of convolution, batch normalization^60^, and ReLU^61^ activations to increase the model’s capability to learn nonlinear functions. We trained the network for 200 epochs using the Adam optimizer^62^ (learning rate = 10^−3^) and batch size of 64 to minimize mean-squared error (MSE) loss. We decayed the learning rate to 10^−4^ at epoch 50 to further improve convergence. The network was trained on a single NVIDIA V100 GPU.

### MorphNet architecture

MorphNet consists of two classes of deep neural networks called variational autoencoder (VAE)^28,29^ and generative adversarial network (GAN)^30,31,37^. The VAE model consists of an encoder and decoder^28^, where the encoder takes raw gene expression count data and learns a low-dimensional latent vector with a Gaussian prior, and the decoder maps the latent representation to parameters of a generative distribution^63^ for each gene in each cell^29^. All VAE architectures used a latent dimensionality of 10 for each dataset. A key advantage of learning cell representations with VAEs is that the latent embeddings are probabilistic, allowing many slightly different embeddings to be sampled for the same cell. This creates an essentially unlimited supply of gene expression and morphology pairs, which helps prevent overfitting.

The generative module of MorphNet is modeled after StyleGAN2^33^, which consists of two deep neural networks called a generator and a discriminator^30^. The generator is responsible for creating morphological images conditioned^37,49^ on latent gene expression, while the discriminator^64^ is responsible for classifying a given image as real or fake. The generator and discriminator are jointly trained^30,65,66^ so that classification scores from the discriminator act as feedback for the generator to improve generated image quality as training progresses. A key feature of StyleGAN2^33^ is its style-based generator^33,34^, which splits the generator into a mapping network and synthesis network (**Supplementary Figure 15a**). The mapping network is a simple 2-layer multilayer perceptron that non-linearly transforms the latent gene expression and random noise into an intermediate latent code^34^, which is incorporated as a “style” into the synthesis network to generate 512 pixels-by-512 pixels RGB morphological images.

### MorphNet Training

We trained the VAE^28,29^ and GAN^30,31^ separately with different loss functions. For VAE^29^, the models were optimized to maximize the evidence lower bound (ELBO)^67^, which encourages the encoder (parameterized by *ϕ*) to learn an approximate posterior distribution *q_ϕ_*(*z*|*x*) given a prior distribution *p_θ_*(*z*) while also encouraging the decoder (parameterized by *θ*) to maximize the likelihood *p_θ_*(*x*|*z*) of the original input being reconstructed^28,67^. Maximizing the ELBO is equivalent to minimizing the following loss:

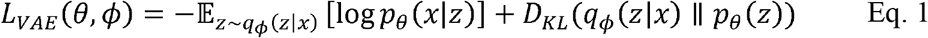

In the above equation, the first term represents the reconstruction loss^28^ for the decoder and the second term represents the distance between the learned posterior distribution from encoder and prior distribution as measured using Kullback-Leibler (KL) divergence^28,67^. All VAE models^28,29^ were trained on a single NVIDIA V100 GPU with batch size of 128 for 100 to 400 epochs depending on dataset using Adam optimizer^62^ with an initial learning rate of 10^−3^ using the Python library scvi-tools^68^.

For GAN, the generator and discriminator were trained to minimize the standard adversarial loss^30^ with various regularizations^66,69^ as described in Karras et al^33^. We used the default hyperparameters recommended in the StyleGAN2^31,33^ paper, except that we changed the regularization strength of R1 regularization^66^ to γ = 10. Briefly, R1 regularization^66^ penalizes the discriminator network gradient on the real data like the gradient penalty used in WGAN-GP^70^ and has been shown to stabilize GAN training^66,71^. The equation for R1 regularization is shown below:

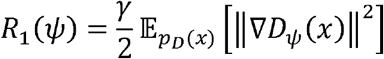

All GAN models^33^ were trained on either NVIDIA V100 GPU or NVIDIA A40 GPU with a batch size of 8 for a maximum of 25 million kimg (number of images seen by the discriminator) using Adam optimizers^62^ (learning rate = 2.5×10^−3^, *β*_1_ = 0, and *β*_2_ – 0.99). We saved the weights of the GAN networks and computed metrics every 200 kimg.

### Augmentation

To train the GAN of MorphNet with limited training data such as Patch-seq, we employed a state-of-the-art augmentation technique for generative models called adaptive discriminator augmentation (ADA)^31^. Previous research has shown that ADA can reduce the number of images required to train a GAN from 10^5^ – 10^6^ to ~10^3^ images^31^. ADA applies a set of image augmentations (pixel blitting, geometric, and color transformations) with probability *p* to both real and generated images before passing them into the discriminator. This is analogous to data augmentation strategiesused to prevent overfitting in image classification tasks. Two key differences are that data augmentation is performed after the generator has produced the images so that the generator doesn’t learn to produce noisy images, and the augmentation probability *p* is adjusted to minimize leakage of augmentation into the generated images (hence the name “adaptive” discriminator augmentation). The value of *p* adapts during training so that *p* is increased/decreased when the discriminator starts to overfit/underfit. Karras et al^31^. proposed a simple heuristic 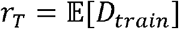, which uses the portion of training set that gets positive discriminator output as a heuristic for discriminator overfitting/underfitting.

## Evaluations

### Fréchet Inception Distance

During training, we periodically evaluated the quality of images using the Fréchet Inception Distance^42^ (FID) metric, which computes the statistical distance between the distribution of generated images to real images. FID is computed by first extracting 2048-dimensional feature vectors for each real and generated image using an ImageNet-pretrained Inception-V3^46,47^ architecture, then computing the Wasserstein-2 distance between the mean and covariance matrices of the feature vectors as shown below:

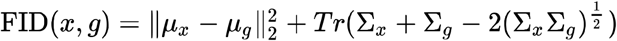

Values of FID can range from zero to infinity, with lower FID score indicating closer distributions between real and generated images and hence better generated image quality^42^. For each dataset, we computed the FID score from 5,000 generated images for each available cell type (i.e. 15K images for BNST, 25K for Isocortex, and 30K for MOp and Patch-seq). We constructed a set of the same number of real images, also balanced by class, by sampling with replacement from the real data, and compared this set of images with the generated images when calculating FID.

### Evaluation by Cell Type Classification

To evaluate whether the generated morphological images reflected the gene expression constraints, we trained a neural network to classify the transcriptional cell type of a morphological image. We finetuned an ImageNet-pretrained ResNet50^43^ model for each dataset, and picked the best classification model based on accuracy and F1 score. After training the ResNet50 classifier on the real images, we used the model to predict transcriptional cell types for the same generated images used in FID calculation (as described in the previous section). An accuracy or F1 score closer to that of the real dataset indicates the generated morphological images reflect gene expression constraints.

To train the classifier, we used an ImageNet-pretrained ResNet50^43^ model with a modified classification module at the end to output the appropriate number of classes. We finetuned the model by freezing the first 7 layers of ResNet50 (Conv, BatchNorm2D, ReLU, MaxPool2D, and three Residual Blocks) and training only the last residual block and the modified classification module. We noticed a significant increase in performance when finetuning the weights associated with the last residual block, which may have enabled the classifier to learn global features uniquely present in morphological images (not found in ImageNet data). Overall, there were a total of 23.8 million parameters, with 15.2 million trainable parameters and 8.5 million non-trainable/frozen parameters. We trained the classifier using standard cross entropy loss using Adam optimizer^62^ with an initial learning rate of 10^−3^, weight decay of 0.0003, and batch size of 64. The classifier was trained on NVIDIA V100 GPU for a total of 200 epochs, with a learning rate scheduler that reduced the learning rate by factor of 10 after every 50 epochs.

### Interpretation of MorphNet

Here we describe methods to interpret the internal representations learned by MorphNet and how groups of specific genes may affect neuronal and nuclear morphology.

### Latent Space Interpolation

To predict potential morphologies of cells transitioning between two cell types (e.g. *Lamp5* to *Vip*), we randomly chose two real cells as the start and end point for the transition. We then obtained the 10-dimensional latent gene expression for each cell using the VAE of MorphNet. Then, we precomputed five evenly spaced (linearly interpolated) intermediate latent positions to use as conditions for the trained MorphNet generator. We repeated this procedure 100 times for *Lamp5* to *Vip* transitions and 100 times for *Pvalb* to *Sst* transitions and used the trained ResNet-50 classifier to predict cell type probabilities for each interpolated step. Note that MorphNet can similarly predict morphology transitions among any two arbitrarily chosen cell types.

### Vector Arithmetic using Differentially Expressed Genes

We used the scanpy^54^ Wilcoxon rank-sum test to identify differentially expressed genes for each cell type in Patch-seq. We considered genes with minimum logfc value of 0.25 and adjusted p-value cutoff below 0.01 to be differentially expressed for each of the six cell types. We then chose two start and end cell types (e.g. *Lamp5* and *Vip*) and subtracted or set equal to zero all differentially expressed genes for the starting cell type. Then, we added values to the differentially expressed genes for the ending cell type, based on the cell’s maximum gene expression value across all genes. This strategy was used to preserve the minimum and maximum raw gene expression value and approximately keep the same library size before and after subtracting and adding values to differentially expressed genes. Finally, from the new raw gene expression vector, we obtained a previously unseen latent gene expression using trained VAE model and fed the latent gene expression to the GAN generator to predict new morphologies.

### Morphological Axes of Variation

We used an unsupervised approach to identify dominant morphological axes learned by MorphNet^72^. Briefly, we performed singular value decomposition (SVD) on the weight matrices learned by the Synthesis network^34^ of the trained MorphNet GAN generator (Supplementary Figure 15**a**). The right singular vectors represent the principal components of the GAN generator weights, which we used to incrementally vary the latent code learned by MorphNet. This helps answer the simple but important question: what specific morphological features can vary (and what are fixed) given a particular gene expression profile? We found that each of the singular vectors represents a distinct morphological feature that can vary for a given gene expression profile. We visualized these morphological features by generating images along two morphological axes for MorphNet trained on MERFISH and MorphNet trained on Patch-seq in Supplementary Figure 15 **b**.

## Data Availability

MERFISH data were downloaded from the Brain Image Library. Patch-seq data were downloaded from https://github.com/berenslab/mini-atlas and Brain Image Library.

## Code Availability

Python implementation of MorphNet is available at https://github.com/single-cell-morphology. The online web tool for predicting neuron morphology is available at https://morphnet.streamlitapp.com/.

## Acknowledgements

We thank Hengshi Yu, Dawen Cai, and Sami Barmada for insightful discussions. This study was supported in part by a National Science Foundation Graduate Research Fellowship to H.L. and by NIH grant RF1MH123199 to J.D.W.

## Supplementary Figures

**Supplementary Figure 1.**
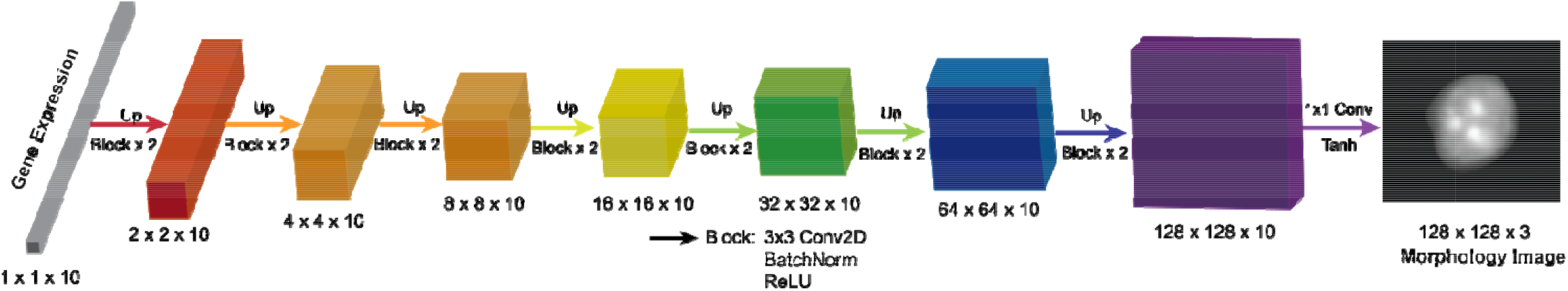
Schematic of CNN Baseline model for predicting single-cell nuclear morphology or neuronal from a latent gene expression vector.

**Supplementary Figure 2.**
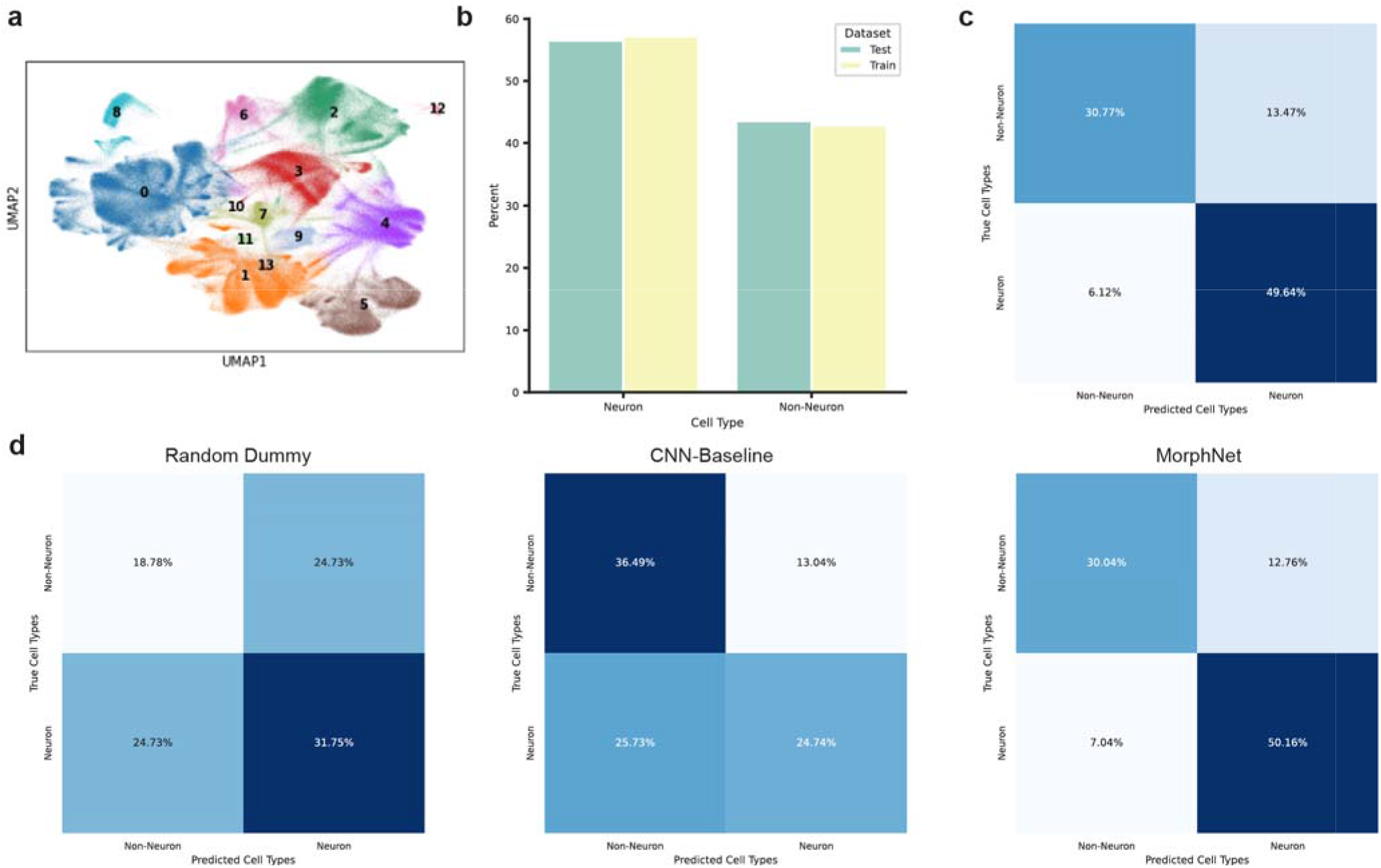
**a.** UMAP of Vizgen MERFISH gene expression data colored by 14 Leiden **clu**sters. **b.** Relative proportions of neuronal and non-neuronal cell types in MERFISH train and test sets. **c.** Con**fu**sion matrix from classifying real MERFISH nuclear morphology images. **d.** (Left) Confusion matrix from classifying real MERFISH nuclear morphology images using Scikit Learn’s Dummy Classifier with “s**tra**tified” strategy. (Middle) Confusion matrix from classifying generated images from baseline model. (Right) C**on**fusion matrix from classifying generated images from MorphNet.

**Supplementary Figure 3.**
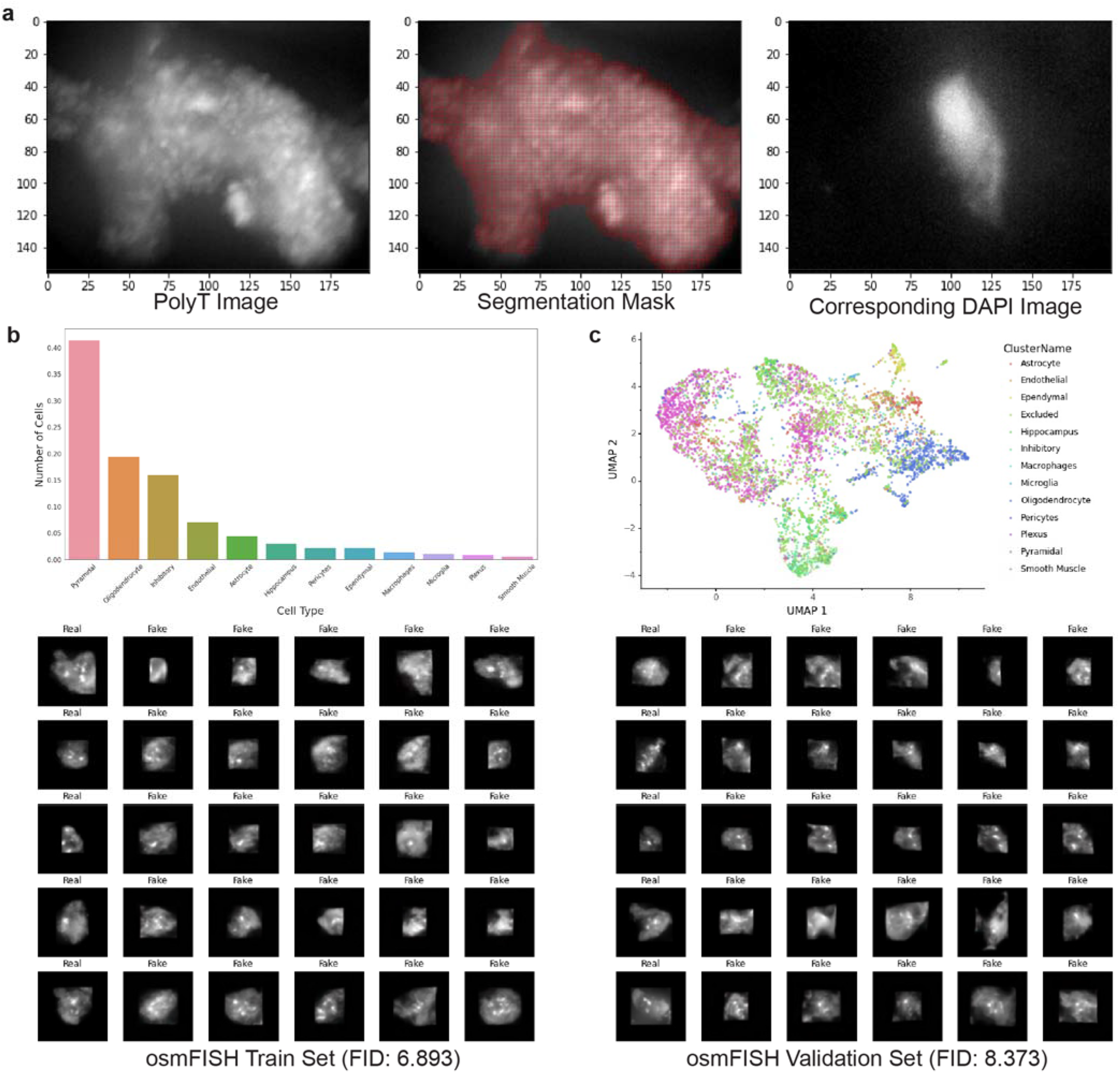
MorphNet results on osmFISH dataset. **a.** To obtain nuclear images from osmFISH, we used the provided segmentation mask image generated based on PolyT staining. Each morphology image spans the size of a single cell, with the nucleus highlighted via DAPI staining. **b.** Cell type distribution of osmFISH. c. UMAP of osmFISH cells clustered using transcriptional (33 genes) signatures of each cell. **d.** Comparison of real and generated images from training set. **e.** Comparison of real and generated images from validation set.

**Supplementary Figure 4.**
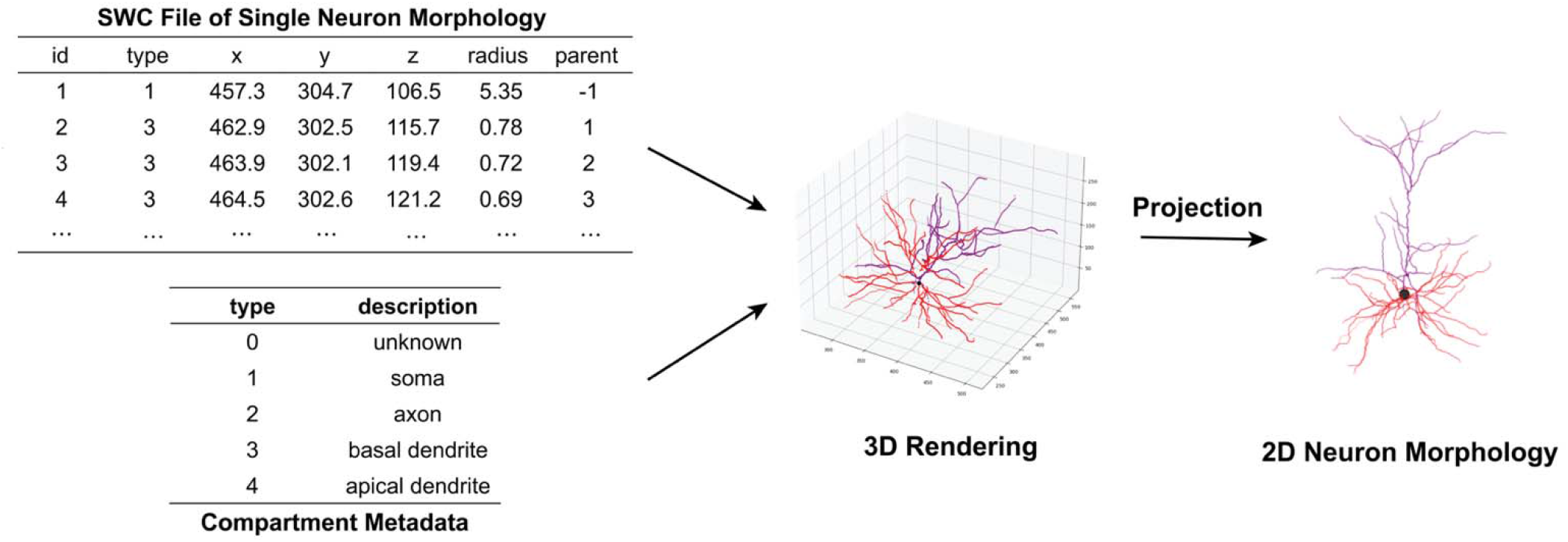
Schematic illustrating the workflow to preprocess raw neuron morphology data into 2-dimensional morphology images to train MorphNet. For each neuron, we projected its 3-dimensional coordinates into the *xy*-plane to create a RGB image of its morphology. Parts of the neurons were colored according to their compartment type.

**Supplementary Figure 5.**
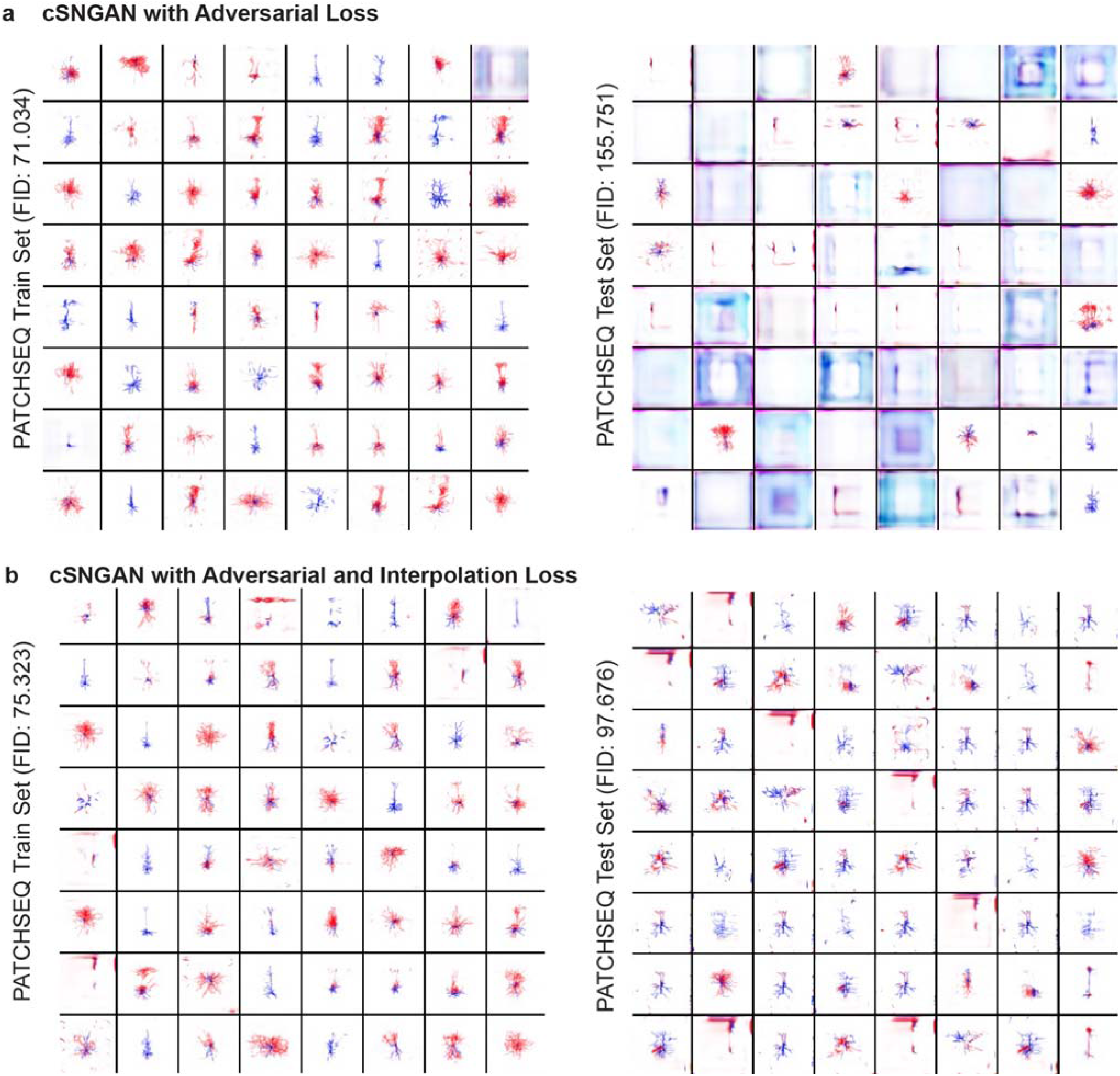
Generated Patch-seq images using cSNGAN. **a.** (Left) Generated morphologies of cells in the Patch-seq training set. (Right) Generated morphologies of cells in the Patch-seq testing set. cSNGAN does not generalize well to test set with just adversarial loss. **b.** (Left) Generated morphologies of cells in the Patch-seq training set using cSNGAN trained with interpolation loss. (Right) Generated morphologies of cells in the Patch-seq testing set using cSNGAN trained with interpolation loss.

**Supplementary Figure 6.**
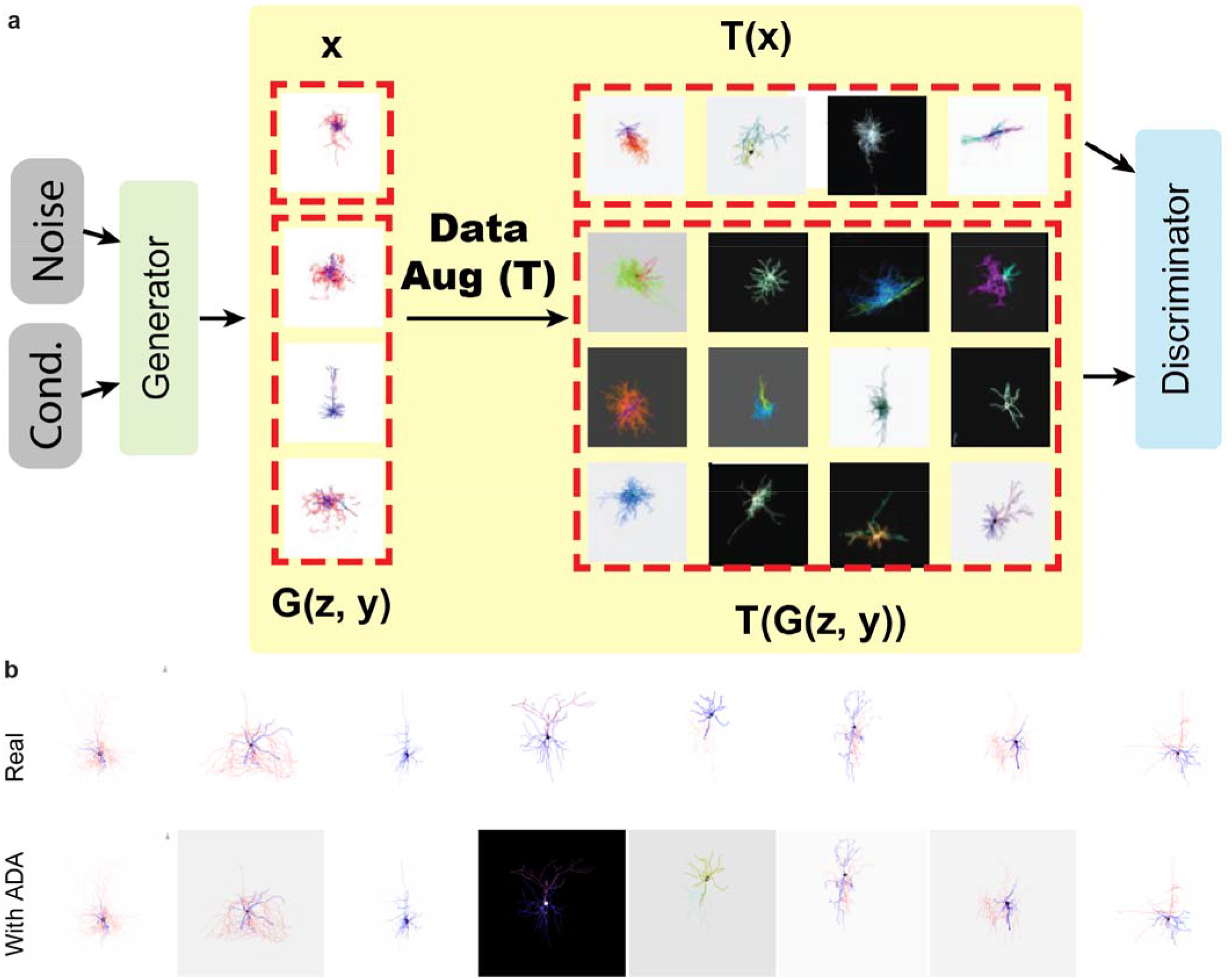
**a.** Schematic of adaptive discriminator augmentation (ADA) technique. ADA applies pixel blitting, geometric, and color transformations to both real and generated images with probability *p* to prevent overfitting the discriminator. **b.** Effect of applying data augmentation to real images with *p* = 0.6 on eight Patch-seq morphology images.

**Supplementary Figure 7.**
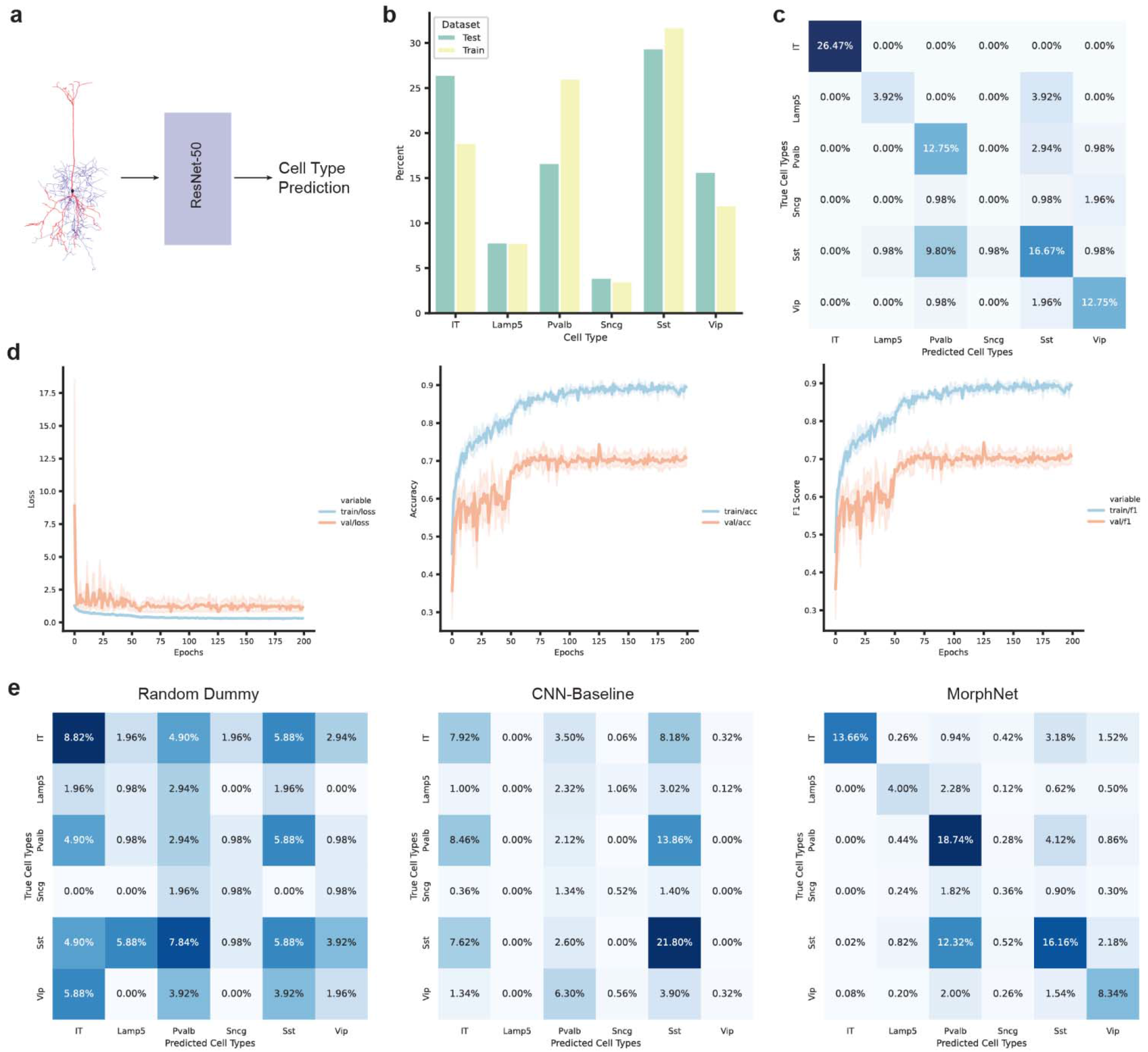
**a.** Schematic of classifying 2D neuron morphologies into one of six cell types (IT, *Lamp5*, *Pvalb*, *Vip*, *Sncg*, or *Sst*). **b.** Cell type distribution between train and test set. Train and test set were split using stratified splitting to match the distribution of cell types as closely as possible. **c.** Confusion matrix of ResNet50-based classifier on test set. **d.** From left to right: Train and validation loss curves, accuracy, and F1 scores over 200 epochs of training. **e.** From left to right: Confusion matrices from dummy classifier, from generated images of CNN-baseline, and from generated images of MorphNet.

**Supplementary Figure 8.**
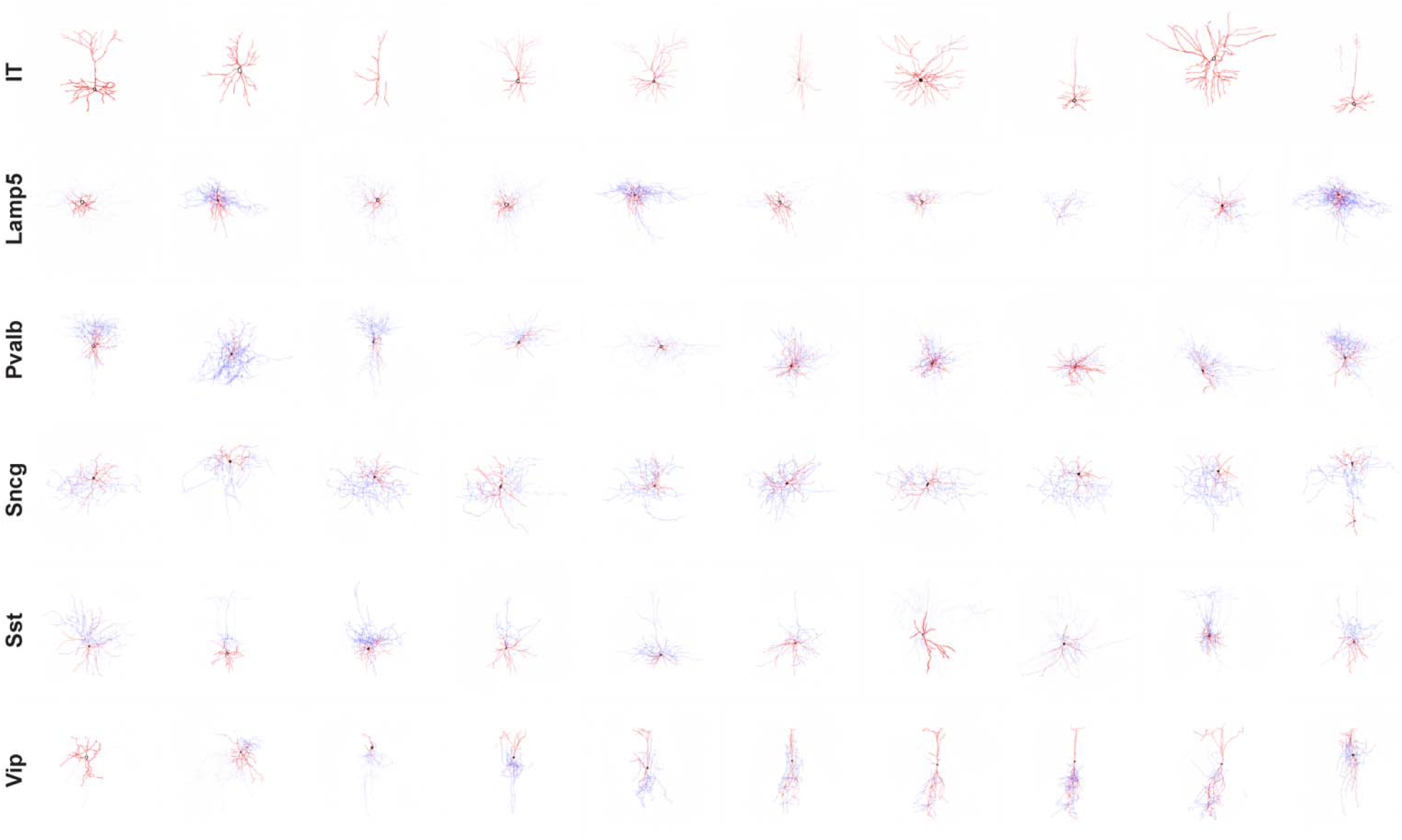
Uncurated 512×512 results generated for MOp dataset for each of the six Patch-seq cell types.

**Supplementary Figure 9.**
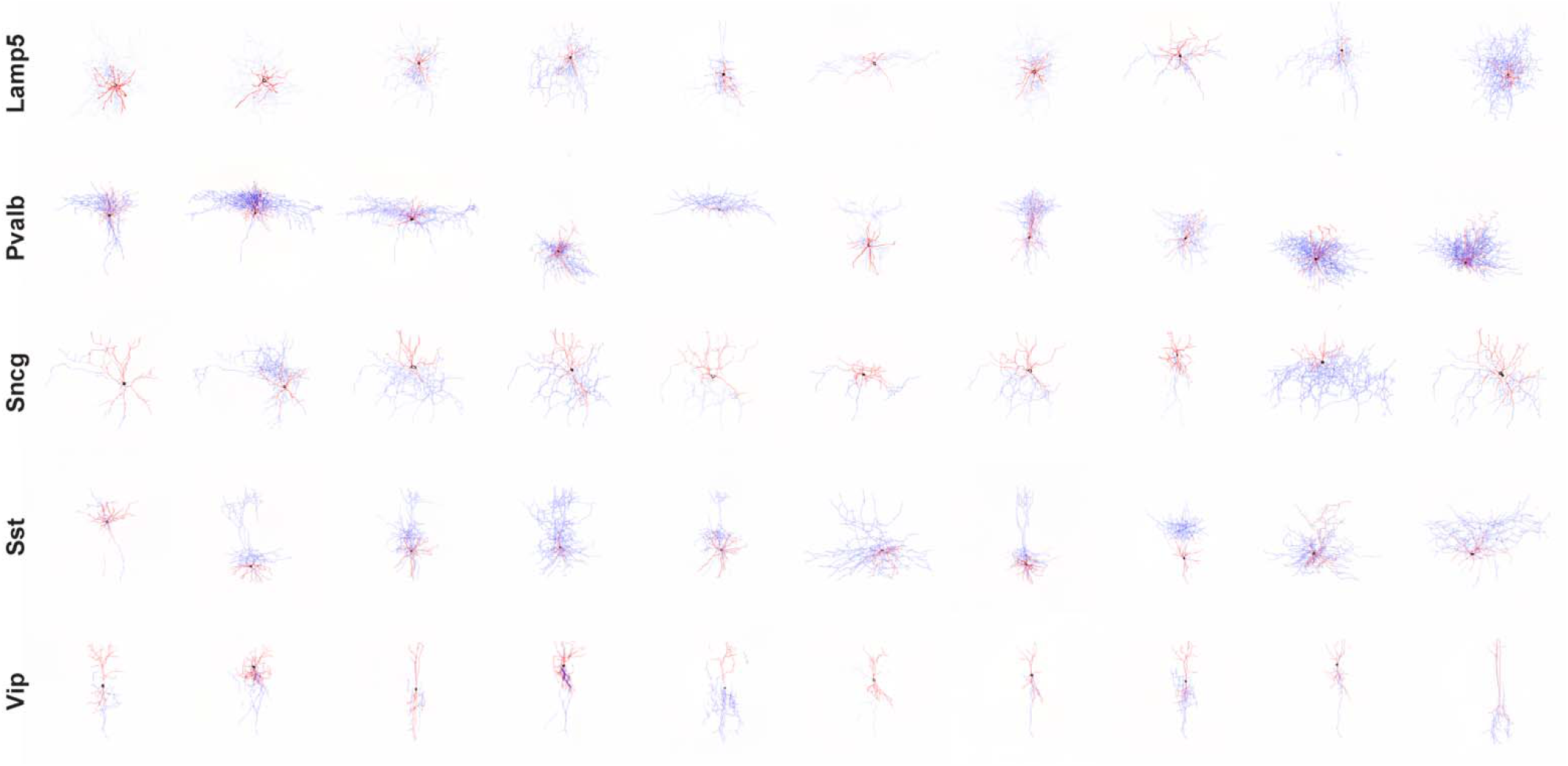
Uncurated 512×512 results generated for Isocortex dataset for each of the five cell types present in the dataset.

**Supplementary Figure 10.**
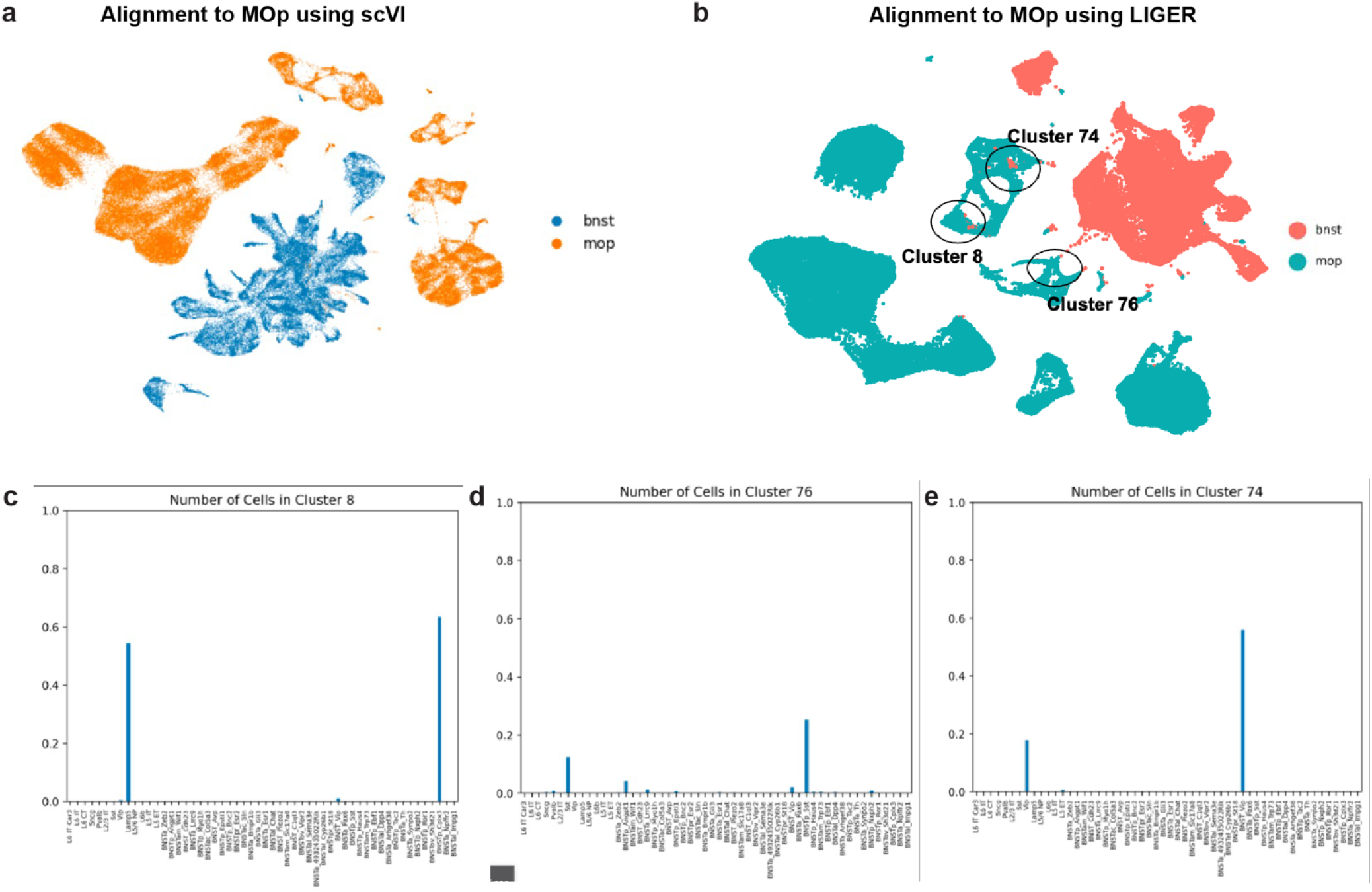
**a.** UMAP plot of BNST and MOP datasets obtained using trained VAE. **b.** UMAP plot of BNST and MOP datasets obtained using LIGER (k=80). **c.** Percentage of cells in Cluster 8, which shows a mixture of *Lamp5* from MOP and BNST_*Cplx3*. **d.** Percentage of cells in Cluster 76, which shows a mixture of *Sst* from MOP and BNST_*Sst*. **e.** Percentage of cells in Cluster 74, which shows a mixture of *Vip* from MOP and BNST_*Vip*.

**Supplementary Figure 11.**
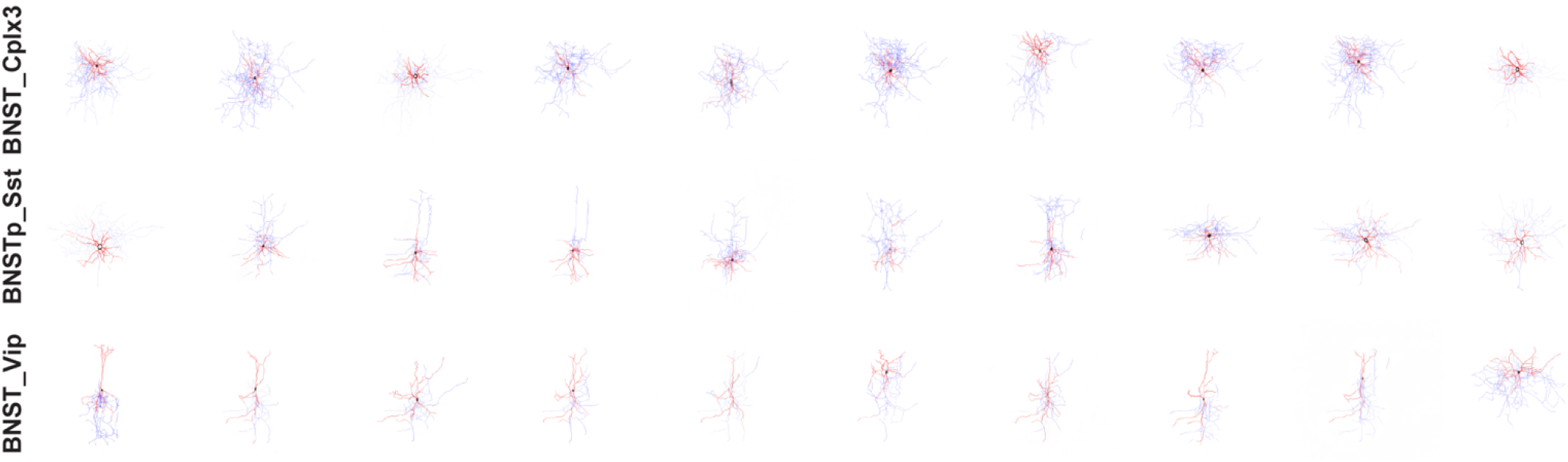
Uncurated 512×512 results generated for BNST dataset for BNST_*Cplx3*, BNSTp_*Sst*, and BNST_ *Vip* clusters.

**Supplementary Figure 12.**
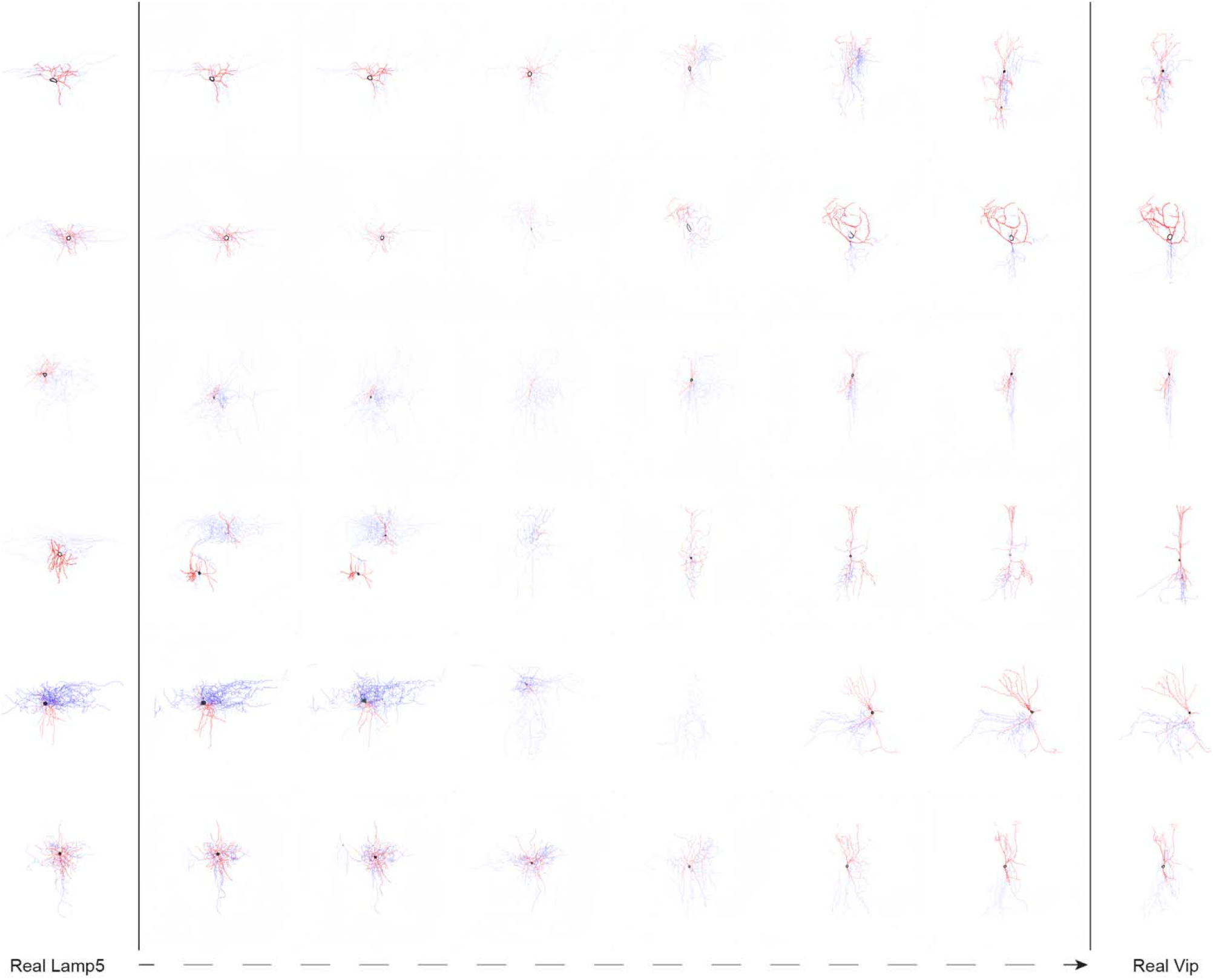
Uncurated 512×512 results for *Lamp5* to *Vip* Transitions.

**Supplementary Figure 13.**
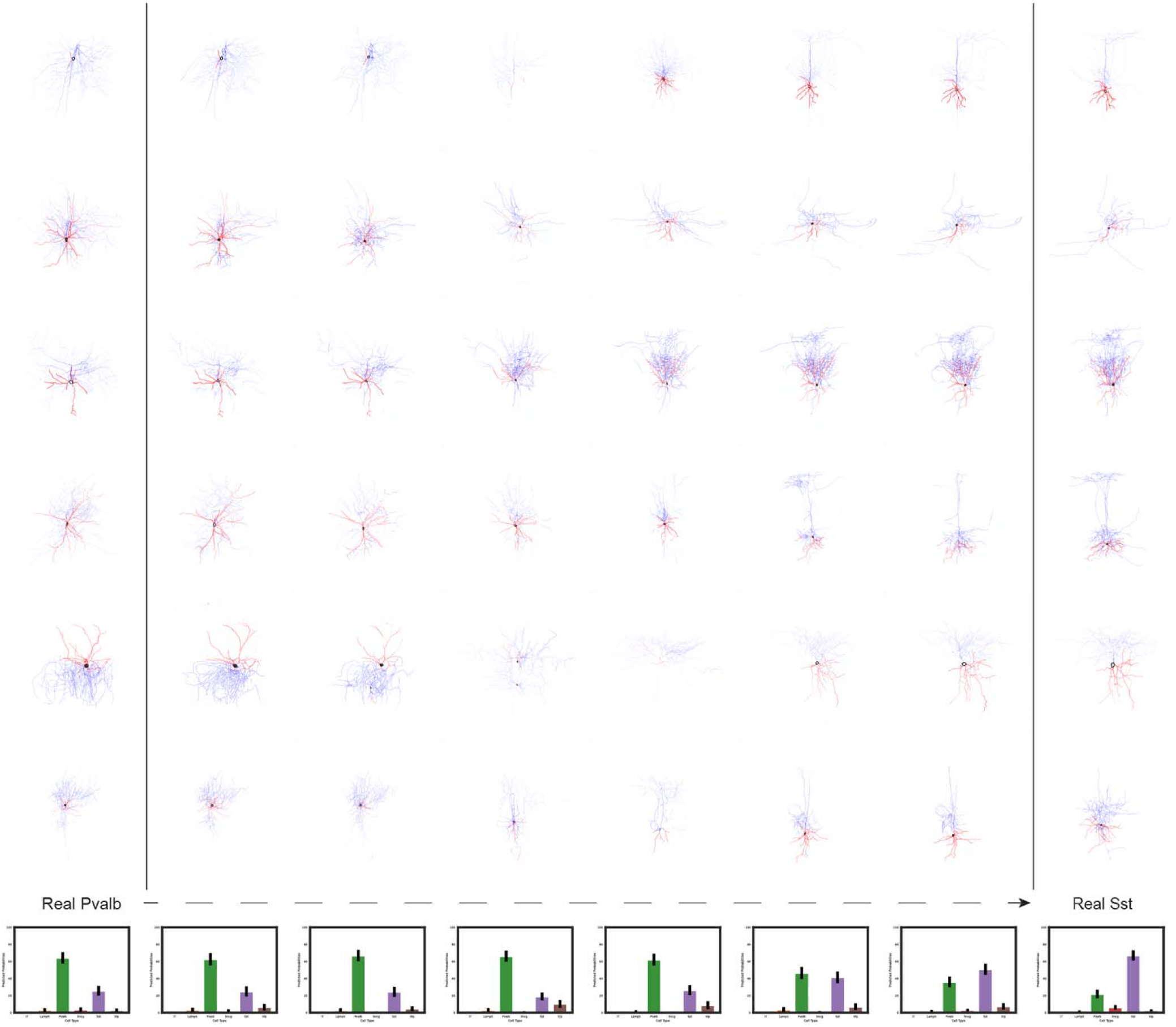
Uncurated 512×512 results for *Pvalb* to *Sst* transitions with cell type probabilities for each step along the transition.

**Supplementary Figure 14.**
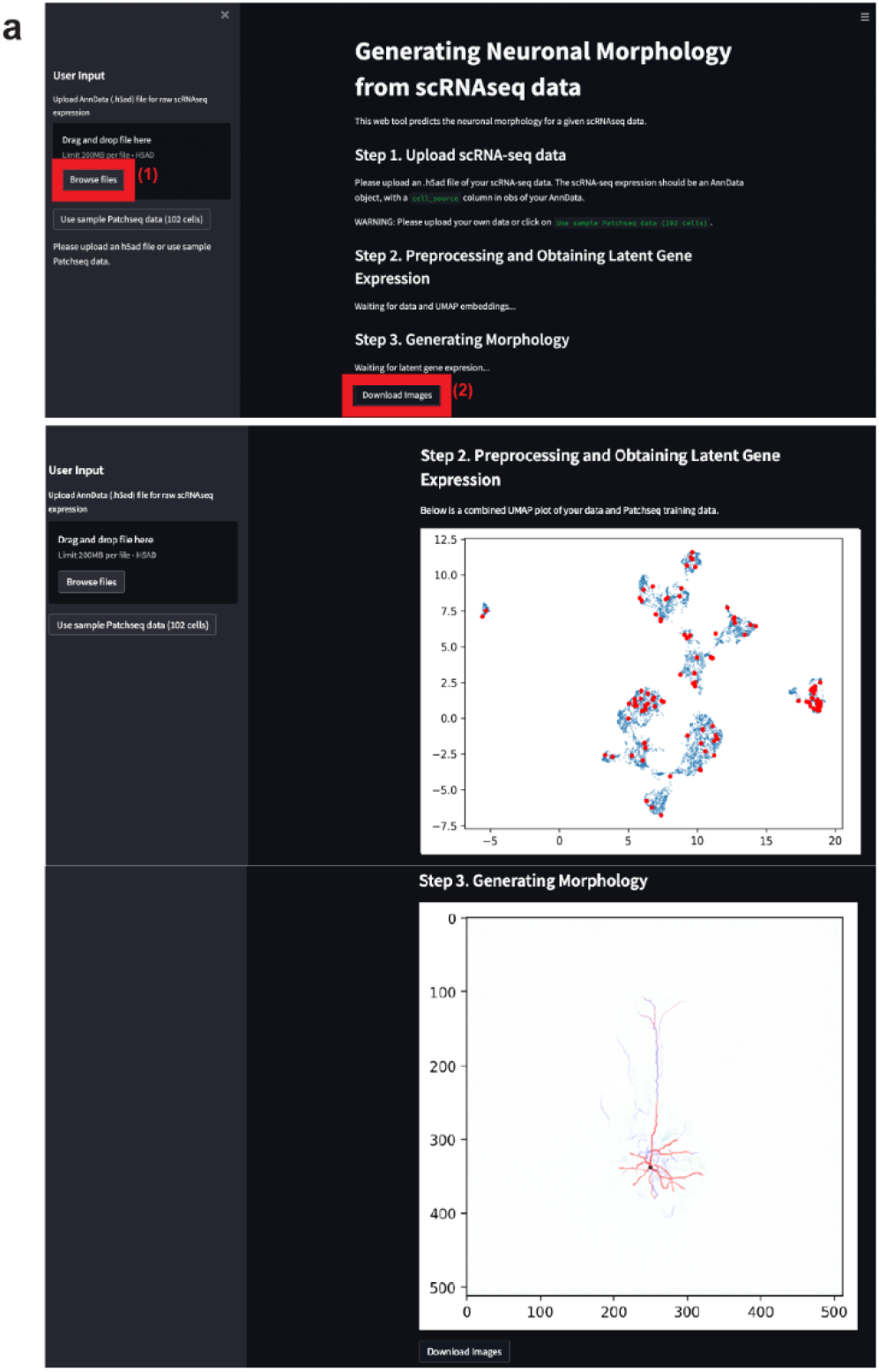
Screenshot of MorphNet webtool created using Streamlit. Users may upload their single-cell RNA-seq dataset in AnnData format (limited to 200 MB per file) or use sample data. Upon uploading the data, the webtool will generate UMAP of the learned latent gene expression and predicted morphologies. Users have the option to download morphology images upon completion.

**Supplementary Figure 15.**
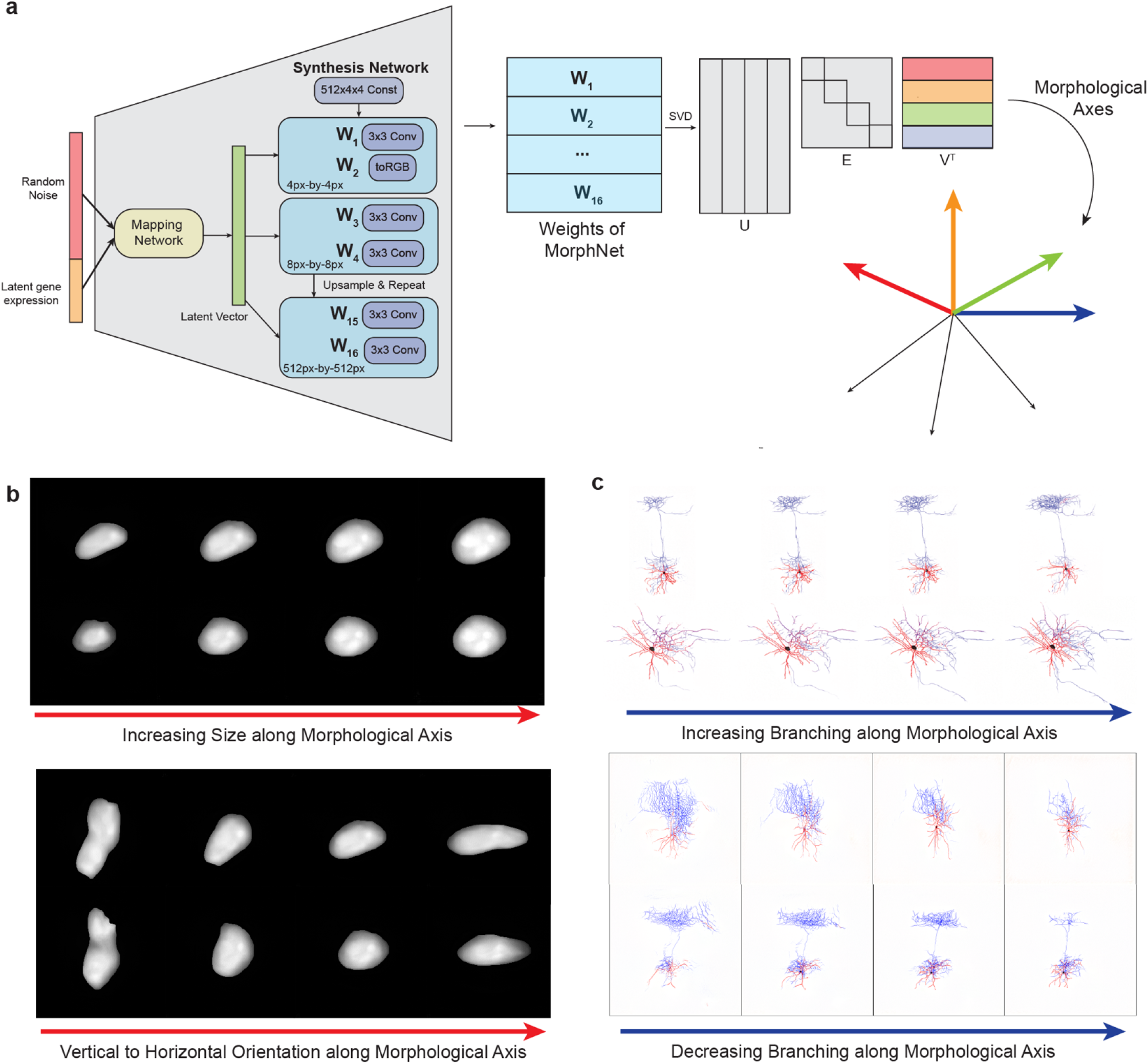
**a.** Schematic for interpretation of learned weights of trained MorphNet using semantic factorization. **b.** Two examples from varying morphological axes for MorphNet trained on MERFISH, which qualitatively affects cell size and vertical/horizontal orientation. **c.** Two examples from varying morphological axes for MorphNet trained on Patch-seq, which qualitatively affects degree of axonal and dendritic branchings.

## References

1. Zink, D., Fischer, A. H. & Nickerson, J. A. Nuclear structure in cancer cells. Nat. Rev. Cancer 4, 677–687 (2004).

2. Minn, A. J. et al. Distinct organ-specific metastatic potential of individual breast cancer cells and primary tumors. J. Clin. Invest. 115, 44–55 (2005).

3. Matsuoka, F. et al. Morphology-Based Prediction of Osteogenic Differentiation Potential of Human Mesenchymal Stem Cells. PLOS ONE 8, e55082 (2013).

4. Li, Q. et al. Leukocyte cells identification and quantitative morphometry based on molecular hyperspectral imaging technology. Comput. Med. Imaging Graph. Off. J. Comput. Med. Imaging Soc. 38, 171–178 (2014).

5. d’Onofrio, G. & Zini, G. Morphology of the Blood. (Taylor & Francis, 1998).

6. Ecker, J. R. et al. The BRAIN Initiative Cell Census Consortium: Lessons Learned toward Generating a Comprehensive Brain Cell Atlas. Neuron 96, 542–557 (2017).

7. Mukamel, E. A. & Ngai, J. Perspectives on defining cell types in the brain. Curr. Opin. Neurobiol. 56, 61–68 (2019).

8. Svensson, V., Vento-Tormo, R. & Teichmann, S. A. Exponential scaling of single-cell RNA-seq in the past decade. Nat. Protoc. 13, 599–604 (2018).

9. Zheng, G. X. Y. et al. Massively parallel digital transcriptional profiling of single cells. Nat. Commun. 8, 14049 (2017).

10. Zhang, M. et al. Molecular, spatial and projection diversity of neurons in primary motor cortex revealed by in situ single-cell transcriptomics. bioRxiv (2020) doi:10.1101/2020.06.04.105700.

11. Moffitt, J. R. et al. Molecular, spatial, and functional single-cell profiling of the hypothalamic preoptic region. Science 362, eaau5324 (2018).

12. Moffitt, J. R. et al. High-throughput single-cell gene-expression profiling with multiplexed error-robust fluorescence in situ hybridization. Proc. Natl. Acad. Sci. 113, 11046–11051 (2016).

13. Chen, K. H., Boettiger, A. N., Moffitt, J. R., Wang, S. & Zhuang, X. Spatially resolved, highly multiplexed RNA profiling in single cells. Science 348, (2015).

14. Zhu, Q., Shah, S., Dries, R., Cai, L. & Yuan, G.-C. Identification of spatially associated subpopulations by combining scRNAseq and sequential fluorescence in situ hybridization data. Nat. Biotechnol. 36, 1183–1190 (2018).

15. Eng, C.-H. L. et al. Transcriptome-scale super-resolved imaging in tissues by RNA seqFISH+. Nature 568, 235–239 (2019).

16. Takei, Y. et al. Integrated spatial genomics reveals global architecture of single nuclei. Nature 590, 344–350 (2021).

17. Codeluppi, S. et al. Spatial organization of the somatosensory cortex revealed by osmFISH. Nat. Methods 15, 932–935 (2018).

18. Yuan, J., Sheng, J. & Sims, P. A. SCOPE-Seq: a scalable technology for linking live cell imaging and single-cell RNA sequencing. Genome Biol. 19, 227 (2018).

19. Liu, Z. et al. Integrating single-cell RNA-seq and imaging with SCOPE-seq2. bioRxiv (2020) doi:10.1101/2020.06.28.176404.

20. Scala, F. et al. Phenotypic variation within and across transcriptomic cell types in mouse motor cortex. bioRxiv (2020) doi:10.1101/2020.02.03.929158.

21. Cadwell, C. R. et al. Electrophysiological, transcriptomic and morphologic profiling of single neurons using Patch-seq. Nat. Biotechnol. 34, 199–203 (2016).

22. Gouwens, N. W. et al. Integrated Morphoelectric and Transcriptomic Classification of Cortical GABAergic Cells. Cell 183, 935–953.e19 (2020).

23. van den Hurk, M., Erwin, J. A., Yeo, G. W., Gage, F. H. & Bardy, C. Patch-Seq Protocol to Analyze the Electrophysiology, Morphology and Transcriptome of Whole Single Neurons Derived From Human Pluripotent Stem Cells. Front. Mol. Neurosci. 11, 261 (2018).

24. Tripathy, S. J. et al. Transcriptomic correlates of neuron electrophysiological diversity. PLOS Comput. Biol. 13, e1005814 (2017).

25. Bomkamp, C. et al. Transcriptomic correlates of electrophysiological and morphological diversity within and across excitatory and inhibitory neuron classes. PLOS Comput. Biol. 15, e1007113 (2019).

26. Bao, F. et al. Integrative spatial analysis of cell morphologies and transcriptional states with MUSE. Nat. Biotechnol. 40, 1200–1209 (2022).

27. Monjo, T., Koido, M., Nagasawa, S., Suzuki, Y. & Kamatani, Y. Efficient prediction of a spatial transcriptomics profile better characterizes breast cancer tissue sections without costly experimentation. Sci. Rep. 12, 4133 (2022).

28. Kingma, D. P. & Welling, M. Auto-Encoding Variational Bayes. http://arxiv.org/abs/1312.6114 (2014) doi:10.48550/arXiv.1312.6114.

29. Lopez, R., Regier, J., Cole, M. B., Jordan, M. I. & Yosef, N. Deep generative modeling for single-cell transcriptomics. Nat. Methods 15, 1053–1058 (2018).

30. Goodfellow, I. J. et al. Generative Adversarial Networks. ArXiv14062661 Cs Stat (2014).

31. Karras, T. et al. Training Generative Adversarial Networks with Limited Data. ArXiv200606676 Cs Stat(2020).

32. Scala, F. et al. Phenotypic variation of transcriptomic cell types in mouse motor cortex. Nature (2020) doi:10.1038/s41586-020-2907-3.

33. Karras, T. et al. Analyzing and Improving the Image Quality of StyleGAN. ArXiv191204958 Cs Eess Stat (2020).

34. Karras, T., Laine, S. & Aila, T. A Style-Based Generator Architecture for Generative Adversarial Networks. ArXiv181204948 Cs Stat (2019).

35. Gayoso, A. et al. scvi-tools: a library for deep probabilistic analysis of single-cell omics data. bioRxiv2021.04.28.441833 (2021) doi:10.1101/2021.04.28.441833.

36. MichiGAN: sampling from disentangled representations of single-cell data using generative adversarial networks | Genome Biology | Full Text. https://genomebiology.biomedcentral.com/articles/10.1186/s13059-021-02373-4.

37. Mirza, M. & Osindero, S. Conditional Generative Adversarial Nets. ArXiv14111784 Cs Stat (2014).

38. Emanuel, G., Moffitt, J. R. & Zhuang, X. High-throughput, image-based screening of pooled genetic-variant libraries. Nat. Methods 14, 1159–1162 (2017).

39. Emanuel, G. & He, J. Using MERSCOPE to Generate a Cell Atlas of the Mouse Brain that Includes Lowly Expressed Genes. Microsc. Today 29, 16–19 (2021).

40. Fuzik, J. et al. Integration of electrophysiological recordings with single-cell RNA-seq data identifies neuronal subtypes. Nat. Biotechnol. 34, 175–183 (2016).

41. Lipovsek, M. et al. Patch-seq: Past, Present, and Future. J. Neurosci. 41, 937–946 (2021).

42. Heusel, M., Ramsauer, H., Unterthiner, T., Nessler, B. & Hochreiter, S. GANs Trained by a Two Time-Scale Update Rule Converge to a Local Nash Equilibrium. ArXiv170608500 Cs Stat (2018).

43. He, K., Zhang, X., Ren, S. & Sun, J. Deep Residual Learning for Image Recognition. ArXiv151203385 Cs (2015).

44. Krizhevsky, A., Sutskever, I. & Hinton, G. E. ImageNet classification with deep convolutional neural networks. Commun. ACM 60, 84–90 (2012).

45. Simonyan, K. & Zisserman, A. Very Deep Convolutional Networks for Large-Scale Image Recognition. ArXiv14091556 Cs (2015).

46. Szegedy, C., Vanhoucke, V., Ioffe, S., Shlens, J. & Wojna, Z. Rethinking the Inception Architecture for Computer Vision. ArXiv151200567 Cs (2015).

47. Szegedy, C. et al. Going Deeper with Convolutions. ArXiv14094842 Cs (2014).

48. Ronneberger, O., Fischer, P. & Brox, T. U-Net: Convolutional Networks for Biomedical Image Segmentation. ArXiv150504597 Cs (2015).

49. Miyato, T. & Koyama, M. cGANs with Projection Discriminator. ArXiv180205637 Cs Stat (2018).

50. Kozareva, V. et al. A transcriptomic atlas of mouse cerebellar cortex comprehensively defines cell types. Nature 598, 214–219 (2021).

51. Tasic, B. et al. Shared and distinct transcriptomic cell types across neocortical areas. Nature 563, 72–78 (2018).

52. Welch, J. D. et al. Single-Cell Multi-omic Integration Compares and Contrasts Features of Brain Cell Identity. Cell 177, 1873–1887.e17 (2019).

53. Saunders, A. et al. Molecular Diversity and Specializations among the Cells of the Adult Mouse Brain. Cell 174, 1015–1030.e16 (2018).

54. Wolf, F. A., Angerer, P. & Theis, F. J. SCANPY: large-scale single-cell gene expression data analysis. Genome Biol. 19, 15 (2018).

55. Cannon, R. C., Turner, D. A., Pyapali, G. K. & Wheal, H. V. An on-line archive of reconstructed hippocampal neurons. J. Neurosci. Methods 84, 49–54 (1998).

56. Stockley, E. W., Cole, H. M., Brown, A. D. & Wheal, H. V. A system for quantitative morphological measurement and electronic modelling of neurons: three-dimensional reconstruction. J. Neurosci. Methods 47, 39–51 (1993).

57. Palacios, J. et al.BlueBrain/NeuroM: v3.2.0. (2022) doi:10.5281/ZENODO.6524037.

58. Yao, Z. et al. A transcriptomic and epigenomic cell atlas of the mouse primary motor cortex. Nature 598, 103–110 (2021).

59. Dumoulin, V. & Visin, F. A guide to convolution arithmetic for deep learning. ArXiv160307285 Cs Stat (2018).

60. Ioffe, S. & Szegedy, C. Batch Normalization: Accelerating Deep Network Training by Reducing Internal Covariate Shift. ArXiv150203167 Cs (2015).

61. Nair, V. & Hinton, G. E. Rectified linear units improve restricted boltzmann machines. in Proceedings of the 27 th International Conference on International Conference on Machine Learning 807–814 (Omnipress, 2010).

62. Kingma, D. P. & Ba, J. Adam: A Method for Stochastic Optimization. ArXiv14126980 Cs (2017).

63. Grün, D., Kester, L. & van Oudenaarden, A. Validation of noise models for single-cell transcriptomics. Nat. Methods 11, 637–640 (2014).

64. Karras, T., Aila, T., Laine, S. & Lehtinen, J. Progressive Growing of GANs for Improved Quality, Stability, and Variation. ArXiv171010196 Cs Stat (2018).

65. Arjovsky, M. & Bottou, L. Towards Principled Methods for Training Generative Adversarial Networks. ArXiv170104862 Cs Stat (2017).

66. Mescheder, L., Geiger, A. & Nowozin, S. Which Training Methods for GANs do actually Converge? ArXiv180104406 Cs (2018).

67. Blei, D. M., Kucukelbir, A. & McAuliffe, J. D. Variational Inference: A Review for Statisticians. J. Am. Stat. Assoc. 112, 859–877 (2017).

68. Gayoso, A. et al. A Python library for probabilistic analysis of single-cell omics data. Nat. Biotechnol. 40, 163–166 (2022).

69. Odena, A. et al. Is Generator Conditioning Causally Related to GAN Performance? Preprint at http://arxiv.org/abs/1802.08768 (2018).

70. Gulrajani, I., Ahmed, F., Arjovsky, M., Dumoulin, V. & Courville, A. Improved Training of Wasserstein GANs. ArXiv170400028 Cs Stat (2017).

71. Roth, K., Lucchi, A., Nowozin, S. & Hofmann, T. Stabilizing Training of Generative Adversarial Networks through Regularization. ArXiv170509367 Cs Stat (2017).

72. Shen, Y. & Zhou, B. Closed-Form Factorization of Latent Semantics in GANs. ArXiv200706600 Cs (2021).

